# Spatiotemporal restriction of *FUSCA3* expression by class I BPC promotes ovule development and coordinates embryo and endosperm growth

**DOI:** 10.1101/612408

**Authors:** Jian Wu, Rosanna Petrella, Sebastian Dowhanik, Veronica Gregis, Sonia Gazzarrini

## Abstract

Spatiotemporal regulation of gene expression plays an important role in developmental timing in plants and animals. FUSCA3 regulates the transition between different phases of development by acting as a link between different hormonal pathways in Arabidopsis. However, the mechanisms governing its spatiotemporal expression patterns are poorly understood. Here, we show that *FUS3* is expressed in the chalaza and funiculus of the mature ovule and seed, but is repressed in the embryo sac, integuments and endosperm. *FUS3* repression requires class I BASIC PENTACYSTEINE (BPC) proteins, which directly bind to the *FUS3* locus and restrict its expression pattern. During vegetative and reproductive development, derepression of *FUS3* in *bpc1*/*2* or *pML1:FUS3* misexpression lines results in dwarf plants carrying defective flowers and aborted ovules. Post-fertilization, ectopic *FUS3* expression in the endosperm increases endosperm nuclei proliferation and seed size and delays or arrests embryo development. These phenotypes are rescued in *bpc1*/*2 fus3*-*3*. Lastly, class I BPCs interact with FIS-PRC2 (FERTILIZATION-INDEPENDENT SEED-Polycomb Repressive Complex 2), which represses *FUS3* in the endosperm. We propose that BPC1/2 promotes the transition from reproductive to seed development by repressing *FUS3* in ovule integuments. After fertilization, BPC1/2 and FIS-PRC2 repress FUS3 in the endosperm to coordinate endosperm and embryo growth.

## INTRODUCTION

Plants integrate endogenous and environmental signals to correctly time the expression of developmental genetic programs. During their life cycle, plants transition through three major phases of development: vegetative, reproductive and seed development. These phase transitions are characterized by large changes in gene expression, which depend on the action of conserved epigenetic machineries. Epigenetic changes are flexible and respond to developmental and environmental cues (Mozgova and Henning, 2015; Mozgova et al., 2015).

Reproductive development in seed plants starts with the production of female and male gametes and is followed by fertilization and seed development. During ovule development the maternal sporophytic integuments originate from the chalaza and enclose the female gametophyte (embryo sac), which contains two gametes: the haploid egg cell and the diploid central cell. The funiculus, connects the ovule to the placental region in the carpel (Drews and Koltunow, 2011; Gasser and Skinner, 2019). After fertilization of the central cell, the triploid endosperm nuclei undergo multiple rounds of division, which are followed by cellularization. In most Angiosperms the function of the endosperm is to nourish the developing embryo. Fertilization of the egg cell generates the diploid zygote, which divides asymmetrically producing two daughter cell lineages that form the apical embryo proper and the basal suspensor, respectively. The integuments will develop into the seed coat only after fertilization of the central cell (Lafon-Placette and Kohler, 2014; Dresselhaus et al., 2016; Gasser and Skinner, 2019). Auxin is a major player in establishing apical-basal polarity and patterning of the embryo, as well as regulating integuments and endosperm development (Figueiredo et al., 2015; Figueiredo et al., 2016; Robert et al., 2018; Lau et al., 2012; de Vries and Weijers, 2017). In the absence of fertilization, seed development is repressed by the Polycomb-Repressive Complex2 (PRC2). In particular, the FIS-PRC2 complex represses autonomous endosperm development, while EMF-PRC2 and VRN-PRC2 prevent seed coat development prior to fertilization (Roszak and Kohler, 2011; Figueiredo and Kohler, 2018).

Seed maturation is characterized by cell expansion and very little cell division. During this stage of development the embryo accumulates storage compounds, acquires dormancy and establishes desiccation tolerance. These processes are largely controlled by the hormone abscisic acid (ABA), the B3 domain family of transcription factors, namely LEAFY COTYLEDON2 <u>(L</u>EC2), ABSCISIC ACID INSENSTIVE3 (<u>A</u>BI3) and <u>F</u>USCA3 (FUS3), as well as the NF-YB subunits of the CCAAT-binding complex, <u>L</u>EC1 and LEC1-LIKE, which are collectively called LAFL (Sreenivasulu and Wobus, 2013). Genetic and spatiotemporal expression analyses together with Chromatin immunoprecipitation (ChIP) and transcriptomic studies suggest that these genes play redundant as well as specific roles in promoting seed maturation, while repressing germination and vegetative development (Sreenivasulu and Wobus, 2013; Jia et al., 2014; Fatihi et al., 2016; Carbonero et al., 2017; Lepiniec et al., 2018). In particular, *FUS3* is a heterochronic gene, which was shown to promote seed maturation by increasing ABA levels while inhibiting vegetative growth and flowering by repressing gibberellins (GA) synthesis (Keith et al., 1994; Curaba et al., 2004; Gazzarrini et al., 2004). These hormones feed back by positively (ABA) and negatively (GA) regulating FUS3 levels (Gazzarrini et al., 2004; Chiu et al., 2016). FUS3 also inhibits vegetative phase change by repressing ethylene signaling (Lumba et al., 2012). Thus, FUS3 regulates phase transitions by modulating hormones syntheses/signaling.

During germination the seed maturation program is repressed by epigenetic mechanisms, which leads to dormancy break and the transition to the next phase of development; these include: CHROMODOMAIN HELICASE DNA BINDING3 (CHD3)/PICKLE (PKL)-dependent chromatin remodeling; Polycomb Repressive Complex2 (PRC2)-mediated histone 3 lysine 27 trimethylation (H3K27me3); H2AK121ub monoubiquitination by the PRC1 components RING-finger homologs AtBMI1A and AtBMI1B; and VIP1/ABI3/LEC (VAL) mediated recruitment of histone deacetylases (HDAC) and PRC complexes (Jia et al., 2014; Lepiniec et al., 2018). Mutations in these genes result in *LAFL* derepression, leading to expression of seed-specific traits and development of embryonic structures in severe mutants. Accordingly, ectopic expression of *LAFL* genes post-embryonically results in similar phenotypes (Lotan et al., 1998; Stone et al., 2001; Gazzarrini et al., 2004; Braybrook et al., 2006). Clearly, multiple pathways ensure a stable repression of the late embryogenesis program during vegetative growth.

Repression of *LAFL* genes has also been observed during early embryonic development. For example, *FUS3* is ectopically expressed in the endosperm of the PRC mutant *medea* (*mea)* (Makarevich et al., 2006), but the mechanism and function of *FUS3* repression in this tissue is unknown. *LAFL* expression is also regulated by post-transcriptional gene silencing; mutants that affect miRNA biogenesis show de-repression of *LAFL* genes in seedlings and early globular stage embryos (Vashisht and Nodine, 2014). This suggests that *LAFL* expression is tightly controlled and subjected to post-transcriptional and epigenetic regulation not only during vegetative growth, but also in specific seed tissues, although the regulation and role of *LAFL* expression during early embryogenesis is far from being fully understood.

*FUS3* transcripts and protein are found as early as the globular stage embryo and become progressively restricted to the protoderm, root and cotyledon tips during mid-embryogenesis (Gazzarrini et al., 2004; Tsuchiya et al., 2004); however, its function during early embryogenesis is unknown. Recently, we have shown that FUS3 plays a critical role also in reproductive development. The *fus3*-*3* loss-of-function mutant displays seed abortion, which is enhanced in plants grown at elevated temperature and dependent on FUS3 phosphorylation (Chan et al., 2017; Tsai and Gazzarrini, 2012). Interestingly, *pML1:FUS3*-*GFP* plants that mis-express *FUS3* during reproductive development also show aborted siliques, suggesting that spatiotemporal expression of *FUS3* must be tightly regulated at this stage of development (Gazzarrini et al., 2004).

To further investigate the role of FUS3 in reproductive development, we have characterized its localization pattern before and after fertilization. Prior to fertilization FUS3 is transiently localized to the integuments and later confined to the chalaza and funiculus of mature ovules, while post-fertilization FUS3 localizes to the seed coat, chalaza and funiculus, aside from the already known localization in the embryo. We show that class I BASIC PENTACYSTEINE (BPC) proteins interact the FIS PRC2 complex and bind to the *FUS3* chromatin. BPC1/2 repress *FUS3* in the stem, integuments of mature ovules, as well as in the endosperm of developing seeds. *FUS3* misexpression in the *bpc1*-*1 and bpc1*-*1 bpc2 (bpc1*/*2)* mutants reduces plant height, impairs the development of flowers, ovule and endosperm leading to seed abortion or arrested embryogenesis. Similar phenotypes are recapitulated in *pML1:FUS3*-*GFP* misexpression plants. Furthermore, the strong vegetative and reproductive phenotypes of *bpc1*/*2* double mutant can be partially rescued in the *fus3*-*3* background, strongly indicating that they are caused by ectopic *FUS3* expression. We propose that during reproductive development BPC1/2- and PRC2-mediated repression of *FUS3* is necessary for ovule development, while after fertilization *FUS3* repression in the endosperm by BPC1/2 and FIS-PRC2 coordinates endosperm and embryo growth. Hence, correct spatiotemporal expression of *FUS3* is required for the transition from plant reproduction to seed development and from pattern formation to seed maturation.

## RESULTS

### FUS3 localizes to reproductive organs before fertilization and is required for ovule development

The *fus3*-*3* loss-of-function mutant displays seed abortion, which is enhanced at elevated temperature (Chan et al., 2017). To investigate the role of *FUS3* in reproductive development, we first determined FUS3 localization pattern in flower buds using a *pFUS3:FUS3*-*GFP* translational reporter (Gazzarrini et al., 2004). However, no FUS3-GFP fluorescence was detected, likely due to the fast turnover rate of FUS3 (Lu et al., 2010). We then used a *pFUS3:FUS3ΔC*-*GFP* reporter, which lacks the PEST instability motif of FUS3 and allows detection of low FUS3 protein levels (Lu et al., 2010). This reporter is non-functional (it doesn’t rescue *fus3*-*3*), but recapitulates *FUS3* expression patterns determined by qRT-PCR, *pFUS3:GUS* and *pFUS3:GFP* reporters (Lu et al., 2010). Using the *pFUS3:FUS3ΔC*-*GFP* reporter, the FUS3 protein was found to be localized to the pistil (septum, valves and funiculus) and ovules, in agreement with microarray data (Figure 1 A-F and Supplemental Figure 1A). In developing ovules FUS3ΔC-GFP was localized to the epidermis of the nucellus, the chalaza, and funiculus, while in mature ovules (FS12) it was localized to the chalaza and funiculus (Figure 1 C-F). After fertilization (6-48 hours after fertilization; HAF) FUS3ΔC-GFP was present in the funiculus, outer layer of the seed coat, chalaza and micropile; it was also localized to the embryo at early stages of embryogenesis (Figure 1G-L and Supplemental Figure 1B).

**Figure 1.**
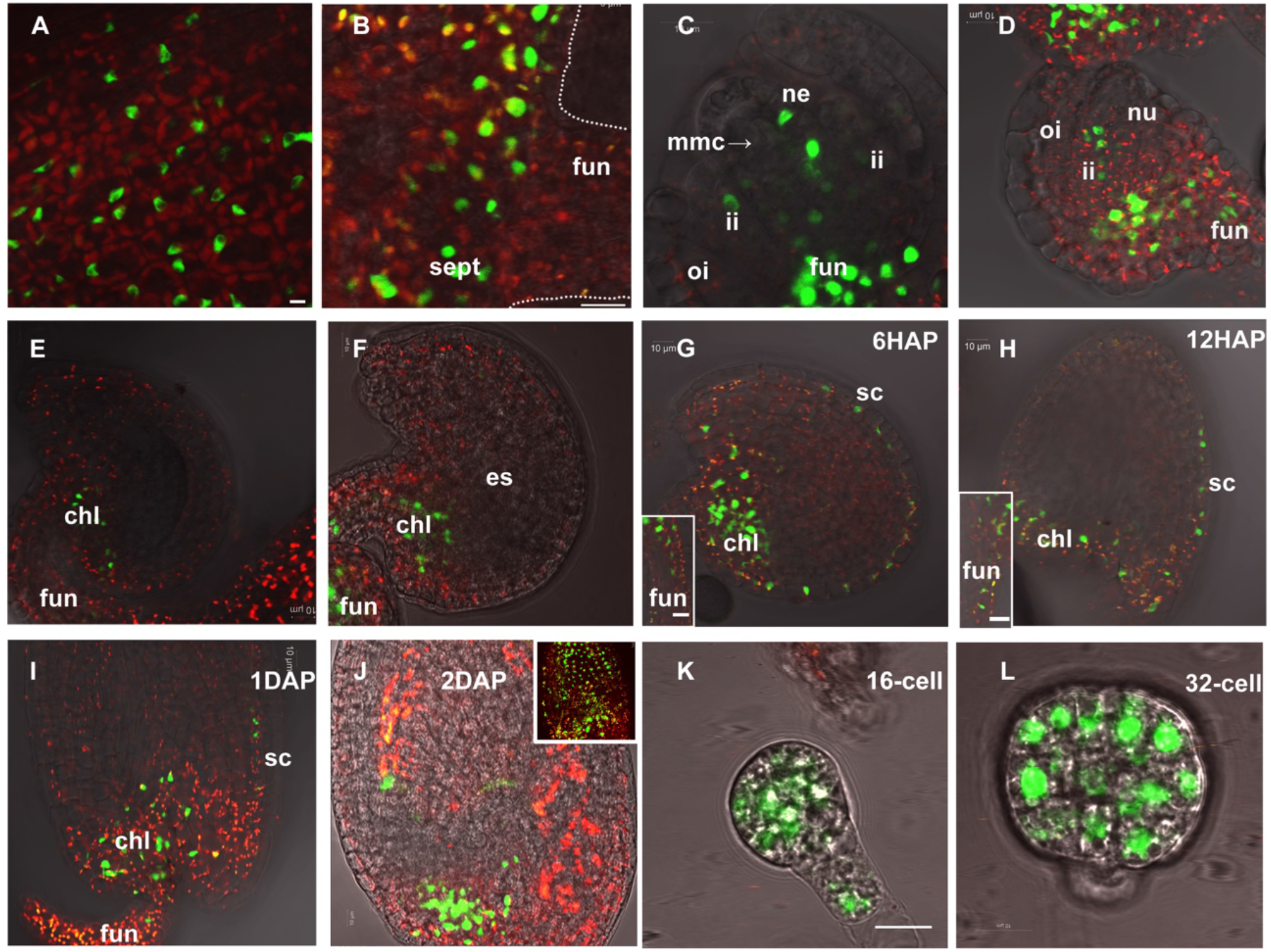
FUS3 localization in developing ovules and during early stages of seed development. Confocal images showing pFUS3:FUS3ΔC-GFP localization in Arabidopsis. **(A)** Valve and **(B)** septum of the pistil. **(C-F)** Developing ovules during female megasporogenesis (**C**) and megagametogenesis at stages FG1-FG7 (**D-F**). FUS3ΔC-GFP fluorescence was localized to the nucellar epidermis (**C**), inner and outer integuments (**C,D**), funiculus, chalazal (**C,F**). (**G-J**) seeds at 6 hours to 2 days (6HAP to 2DAP) after pollination. FUS3ΔC-GFP fluorescence was localized to the seed coat, chalaza and funiculus (**G-J**). **(K)** Suspensor and 16-cell stage embryo proper. (**L**) 32-cell stage embryo proper. chl: chalaza; es, embryo sac; fun, funiculus; ii: inner integument; megaspore mother cell; ne, nucellar epidermis; nu: nucellus; oi: outer integument; sept, septum. Red, autofluorescence from chlorophyll. Purple dashed lines represent the outline of embryo sac. Scale bars, 10μm.

To further address the role of *FUS3* in reproduction, we monitored ovule development in *fus3*-*3* loss-of-function mutant and *pML1:FUS3*-*GFP* misexpression lines (Gazzarrini et al., 2004). *pML1:FUS3*-*GFP* was shown to rescue all *fus3*-*3* seed maturation defects, including desiccation intolerance, however misexpression during postembryonic development caused additional phenotypes (Gazzarrini et al., 2004). Strong *pML1:FUS3*-*GFP* lines show delayed vegetative growth and flowering, reduced plant height and aborted siliques, as previously described (Figure 2A; Gazzarrini et al., 2004; Lu et al., 2010). In addition, we found that in intermediate-to-strong *pML1:FUS3*-*GFP* lines FUS3-GFP was mislocalized to the endothelium, outer and inner integuments of developed ovules, while in aborted ovules FUS3-GFP surrounded the aborted embryo sac (Figure 2B). After fertilization, *pML1:FUS3*-*GFP* seeds showed FUS3-GFP mislocalization to the endosperm (Figure 2B). Moreover, by opening the developed siliques of intermediate-to-strong *pML1:FUS3*-*GFP* lines we also found that they contained aborted seeds or seeds with delayed development (Figure 2C, D). To determine if seed abortion in *fus3*-*3* and *pML1:FUS3*-*GFP* is the result of impaired ovule development, we analyzed ovules before fertilization and compared them with wild type (Figure 2E). The embryo sac of wild type ovules at FS12 stage contained the egg nucleus, the secondary endosperm nucleus, the synergids, and was surrounded by inner and outer integuments. However, at FS12 stage the embryo sac of some *fus3*-*3* and *pML1:FUS3-GFP* lines was delayed at various stages, from FG1 to FG6, arrested or not fully wrapped by the integuments (Figure 2E). The arrest of female megagametogenesis resulted in seed abortion in *fus3*-*3* and more so in strong *pML1:FUS3-GFP* lines (Figure 2C, D).

**Figure 2.**
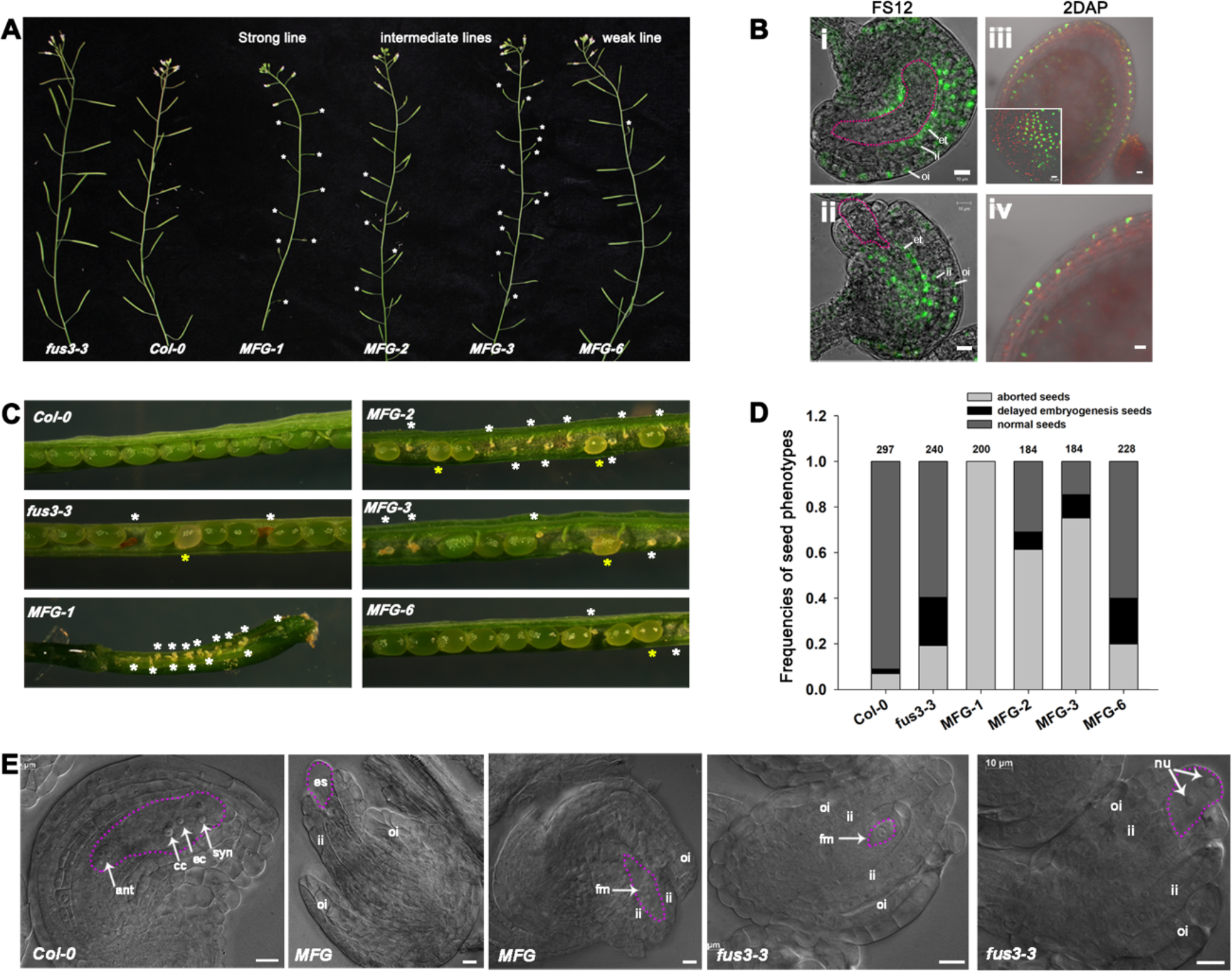
*FUS3* is required for ovule development. **A**, Aborted silique (asterisks) in *fus3*-*3 pML1:FUS3*-*GFP* (*MFG*) overexpression lines. **B**, pML1:FUS3-GFP localization to the integuments and endothelium of ovules at flower stage 12 (FS12), and outer layer of the seed coat and endosperm (inset) of 2DAP seeds. (**i**) developed ovule; (**ii**) aborted embryo sac; (**iii, iv**). outer layer of the seed coat and the endosperm (inset) in 2DAP seeds Bar, 10μM. **C**, Aborted seeds (white asterisk) and delayed embryogenesis (yellow asterisk) in *MFG* and *fus3*-*3* siliques. **D**, The distribution of seeds in peeled, half sides siliques of WT, *MFG* and *fus3*-*3* (n= ten siliques/genotype). **E**, DIC images of WT, *MFG* and *fus3*-*3* FS12 ovules. Pink dashed lines outline the embryo sac. Ant: anti: antipodals; ec: egg cell; es: embryo sac; et: endothelium; fm: functional megaspore; ii, inner integument; nu: nuclei; oi, outer integument; syn: synergid cell nuclei. Bars represent 10μm

Taken together, these results show that spatiotemporal localization of FUS3 is tightly regulated and that lack or misexpression of *FUS3* severely impairs embryo sac and integument development, indicating that spatiotemporal control of *FUS3* expression is required for proper ovule development.

### Class I BPC transcription factors bind to (GA/CT)_n_ motifs in *FUS3*

To understand the mechanisms controlling the spatiotemporal patterns of *FUS3* expression, we identified upstream regulators of *FUS3* by yeast one-hybrid. To increase screening specificity, a short genomic region of 615bp upstream of the *FUS3* translation start (*pFUS3*) was screened against an Arabidopsis transcription factor library (Figure 3A; Mitsuda et al., 2010). About 200,000 yeast transformants were screened and 69 grew on selection plates. Sequencing of the cDNA inserts revealed that all colonies contained BPC3. BPCs are a small group of plant specific transcription factor with six genes and a pseudogene (BPC5) that are divided into 3 classes based on sequence similarity: class I (BPC1/2/3), class II (BPC4/5/6) and class III (BPC7) (Meister et al., 2004). We retested individually all class I BPCs (BPC1-3) and also included class II BPC4, which is not present in the cDNA library but it is highly expressed in embryos and flowers (Berger et al., 2011). The results show that all three class I BPCs bound to *pFUS3* by yeast one-hybrid, but not class II BPC4 (Figure 3A).

**Figure 3.**
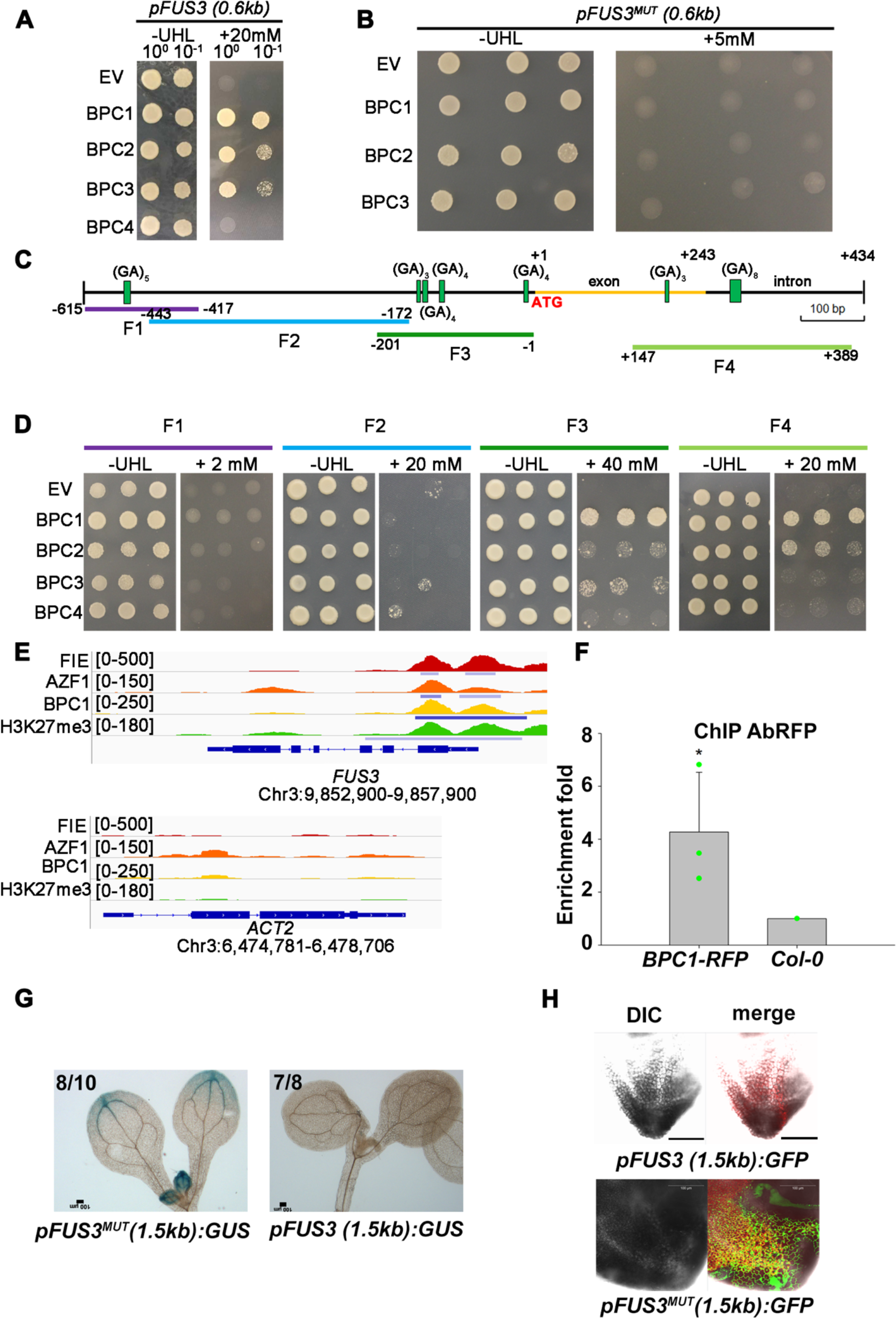
Class I BPCs bind to the *FUS3* genomic region proximal to the transcription start site. **A**, BPC1/2/3 bind to a *FUS3* genomic region of 615bp upstream of the translation start [*pFUS3(0.6 kb)*; −615 to +1 base pairs]. **B**, BPC1/2/3 do not bind the *FUS3* genomic sequence carrying mutations in (GA/CT)_n_ motifs [*pFUS3*^*MUT*^*(0.6 kb)*]. Colonies in **A** and **B** were selected on -ura-his-leu medium (-UHL) with or without 5 or 20mM 3-AT. **C**, Distribution of (GA/CT)_n_ motifs in *FUS3* genomic sequence (−615 to +434). **D**. Binding specificity of BPC1/2/3 to truncated *FUS3* genomic sequences shown in C (F1 to F4). **E**, Bowser view of chromatin occupancy of FIE, BPC1, AZF1 and H3K27me3 at *FUS3* and *ACT2* (negative control) in 30-h-old seedlings using ChIP-seq data from Xiao et al. (2017). Numbers indicate peak Significant peaks (Q < 10^−10^) according to MACS2 are marked by horizontal bars. **F**. Real-time PCR analysis of ChIP assay using chromatin extracted from *35S:BPC1*-*RFP* and *Col*-*0* (negative control) inflorescences and primers for the F3 region of *pFUS3*. Antibodies against the RFP tag were used in the IP. Error bars represent the propagated error value using three biological replicates (*: p<0.05; student t-test). **G**, *pFUS3(1.5kb):GUS* and *pFUS3*^*MUT*^*(1.5kb):GUS* stain in 10-days-old seedlings; numbers refer to the number of transgenic lines displaying the same GUS stain pattern as shown in **G**. **H**, *pFUS3(1.5kb): GFP* and *pFUS3*^*MUT*^*(1.5kb):GFP* fluorescence in the leaf tip of 15-days-old seedlings.

BPCs were shown to bind to (GA/CT)_n_ cis elements in several plant species, with a preference for different numbers of repeats (Berger and Dubreucq, 2012; Simonini and Kater, 2014). When all (GA/CT)_n_ motifs of the *pFUS3* were mutated (*pFUS3*^*MUT*^), none of the class I BPCs interacted with the *FUS3* sequence, confirming binding specificity (Figure 3B; Supplemental Figure 2). To identify the binding location of BPCs on *pFUS3*, we generated truncations of approximately 200bp fragments (F1 to F3); the first exon/intron region containing 2 (GA/CT)_n_ repeats (F4) was also tested (Figure 3C). In Y1H, BPC1 showed strong binding whereas BPC2/3 weak binding to the 5’UTR (F3) and first exon/intron regions (F4), where (GA/CT)_n_ motifs are enriched (Figure 3D). BPC1/2/3/4 did not bind the promoter region further upstream, corresponding to the F1 or F2 truncations, where there is only one (GA)_5_ or no (GA/CT)_n_ motif, respectively (Figure 3D). To determine if BPC1 also binds to the *FUS3* locus *in vivo* during reproductive development, we generated BPC1 overexpression lines and performed ChIP in inflorescences, which show that BPC1 binds to this region (Figure 3F).

Altogether, this indicates that class I BPCs bind to the 5’UTR and first intron/exon regions of *FUS3* in Y1H. Furthermore, BPC1 also binds to *FUS3 in vivo* during reproductive development.

### Class I BPCs repress *FUS3* during vegetative growth

In a genome-wide study, BPC1 was found to interact with and recruit the conserved PRC2-complex subunit FIERY (FIE; Supplemental Figure 4) *in vivo* and trigger polycomb-mediated gene silencing in imbibed seeds (Xiao et al., 2017). We first analyzed ChIP-seq data from Xiao et al. (2017) and found that the first exon/intron and 5’UTR of *FUS3* was bound by BPC1, but not the ACTIN (ACT2) control, in seedlings (Figure 3E). Furthermore, this same region was bound by FIE and associated with H3K27me3, a repressive mark (Figure 3E). Lastly, BPC1/2 interact with EMBRYONIC FLOWER2 (EMF2), which belongs to the EMF-PRC2 complex involved in repressing the vegetative-to-reproductive and embryo-to-seedling phase transitions (Supplemental Figure 4; Xiao et al., 2017; Mozgova et al., 2015). This suggests that *FUS3* may be repressed in germinating seeds by BPC1 recruitment of EMF-PRC2. To confirm this, we mutated all BPC binding sites (GA/CT)_n_ in the *FUS3* sequence (*pFUS3*^*MUT*^) and showed that *pFUS3*^*MUT*^*:GUS/GFP* is indeed derepressed post-embryonically in leaves and root tips (Figure 3G,H). Together with previous data showing that *FUS3* was strongly upregulated in *swinger curly leaf* (*swn clf*) (Makarevich et al., 2006), these results strongly suggest that BPC1 binds to and represses *FUS3* during vegetative development by recruiting the EMF-PRC2 complex.

### Class I BPCs form homo- or heterodimers and recruit FIS-PRC2

Previous ChIP assays showed that in closed flowers the *FUS3* locus is also associated with the FIS-PRC2 complex component MEA and H3K27me3 repressive marks, and that *FUS3* is upregulated in the endosperm of *mea*/*MEA* seeds at 3 days after flowering (DAF) (Makarevich et al., 2006). Given that BPC1 bind to the *FUS3* locus in closed flowers (Figure 3F), we hypothesized that *FUS3* may also be repressed during reproductive development by one or more class I BPCs through FIS-PRC2 recruitment. To test this hypothesis, we first determined if all class I BPCs interact *in planta* with the FIS-PRC2 complex, which acts during gametophyte and endosperm development (Figure 4). All class I BPCs interacted with the unique components of this PRC2 complex, FIS2 and MEA, and also with the PRC2-shared component, MSI1, in BiFC assays; all but BPC3 also interacted with FIE (Figure 4). In agreement with previous Y2H results, class I BPCs also interacted with each other *in planta*, and BPC2 and 3 could also form homodimers (Figure S3; Simonini et al., 2012). No class I BPC member or FIS-PRC2 component interacted with FUS3, suggesting that these BiFC interactions are specific (Supplemental Figure 5). Lastly, given that BPC6 recruits PRC2 by interacting with LIKE HETEROCHROMATIN PROTEIN1 (LHP1; Hecker et al., 2015), we also tested the interaction between class I BPCs and LHP1 *in planta*. However, the results showed no interaction among them, suggesting class I and class II BPCs recruitment of the PRC2 complex may differ (Supplemental Figure 6). We conclude that class I BPCs can form homo-and heterodimers and recruit the FIS-PRC2 complex *in planta*.

**Figure 4.**
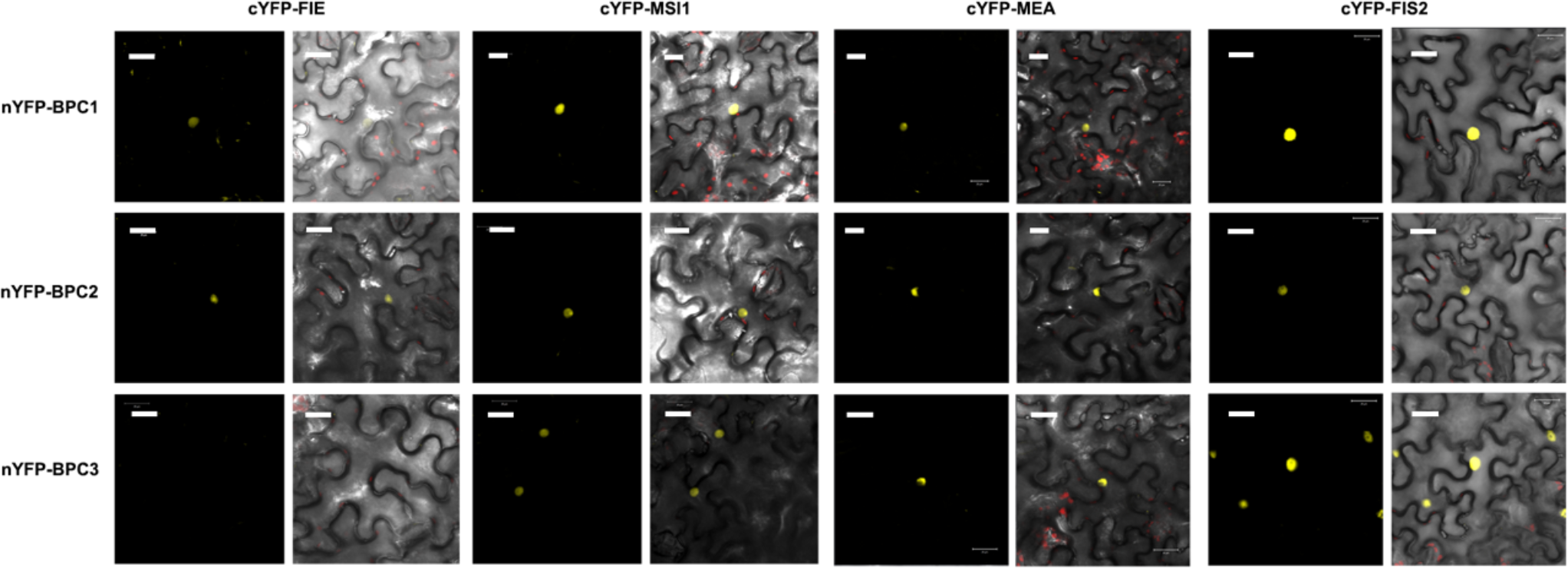
Class I BPC family members intact with FIS-PRC2 complex. The interaction between Class I BPC family members and FIS complex in *N. benthamiana* by Bimolecular Fluorescence Complementation (BiFC). Lack of interaction between FUS3 and BPCs or FIS-PRC2 in BiFC assays is shown as the negative control (Supplemental Figure 5).

Class I BPCs were shown to be expressed in ovules (Monfared et al., 2011). To have a better understanding of the spatiotemporal expression pattern of class I BPCs during reproductive development and embryogenesis, we tracked their expression patterns before (FS4-12) and after (1-11DAF) fertilization using transcriptional or translational reporters. Class I BPCs had largely overlapping expression patterns before fertilization and they were all highly expressed in almost all tissues of developing ovules, while soon after fertilization BPCs were expressed in embryos from the globular to the cotyledon stage, as well as the endosperm and seed coat (Figure 5). BPC1 had a more restricted pattern before (chalaza and micropile) and after (chalaza, micropile, seed coat) fertilization. This suggests that class I BPCs act redundantly during ovule and embryo development. As previously shown, the FIS-PRC2 complex subunits FIS2 and MEA were only expressed in the central cell of developing ovules and in the endosperms at 2DAF (Supplemental Figure 7; Wang et al., 2006).

**Figure 5.**
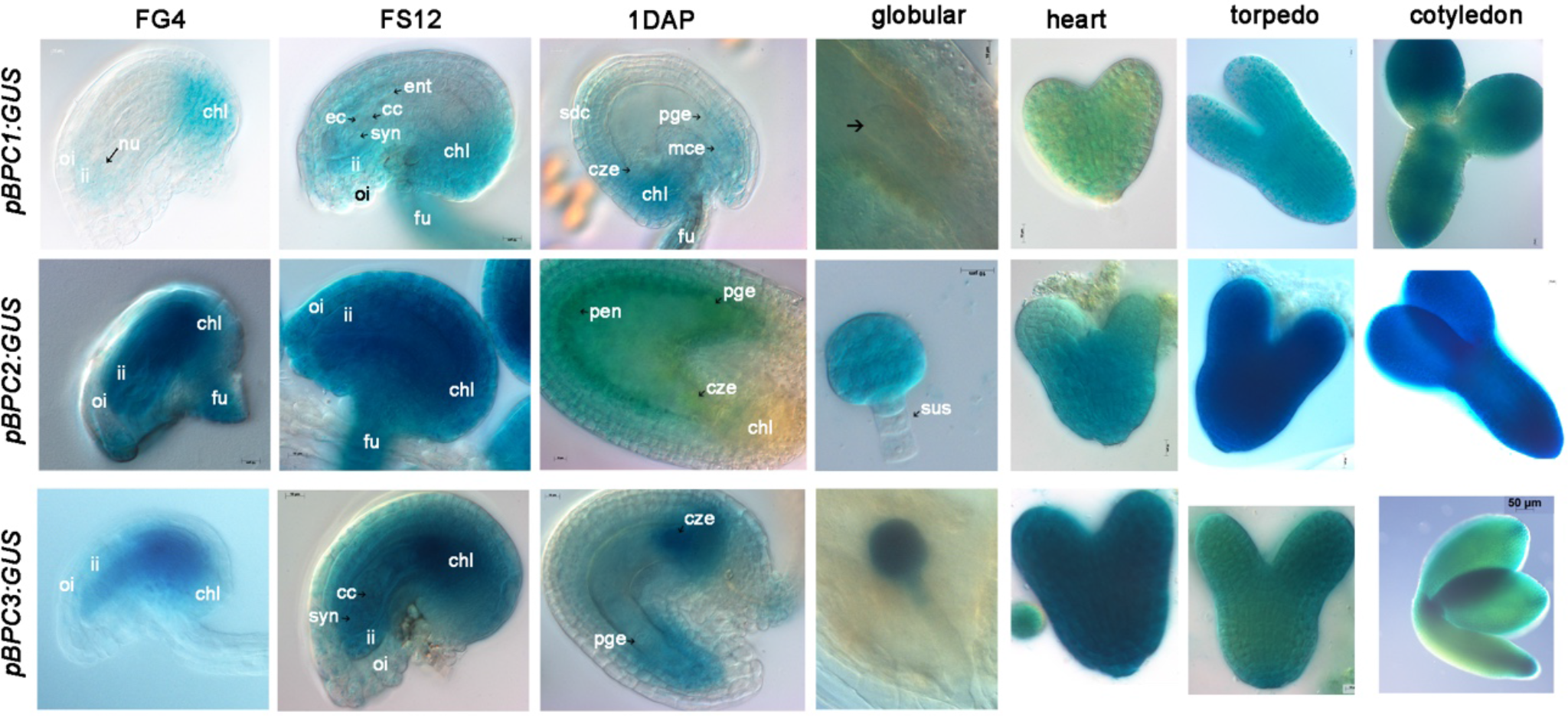
Expression patterns of Class I *BPCs* in ovules and embryos. Class I BPC1s expression patterns in ovules before pollination at flower stages FG4 and FS12; in seeds at 1 day after pollination (DAP); and in embryos at globular, heart, torpedo and cotyledon stages. Ant: antipodals; cc: central cell; chl: chalaza; cze: chalazal endosperm; ec: egg cell; fu: funiculus; ii: inner integuments; mce: micropilar endosperm; oi: outer integuments; pen: peripheral endosperm; pge: pre-globular embryo; sdc: seed coat; sus: suspensor; syn: synergids cell.

These data show that BPCs can interact with each other and with FIS-PRC2 to regulate gene expression. Given the specific localization of FIS and MEA to the central cell and endosperm, and *FUS3* derepression in the endosperm of *mea*/MEA, we conclude that aside from their role in silencing *FUS3* during vegetative growth through EMF-PRC2, class I BPCs repress *FUS3* during reproductive and seed development by recruiting FIS-PRC2 in the central cell and endosperm. Furthermore, BPCs may recruit sporophytic PRC2 (EMF/VRN PRC2) to repress FUS3 in the integuments and seed coat.

### Reproductive defects of *bpc1*/*2* are partially rescued by *fus3*-*3*

Previously, *bpc* mutants were shown to display pleiotropic phenotypes during vegetative and reproductive development (Monfared et al., 2011). Higher order *bpc1*/*2 and bpc1*/*2*/*3* mutants are dwarf, have shorter or aborted siliques, display severe seed abortion and defects in embryo sac development, while most single *bpc* mutants resemble wild type, suggesting functional redundancy (Figure 6A-F; Supplemental Figure 8A-D; (Monfared et al., 2011). These phenotypes are remarkably similar to those shown by *pML1:FUS3* misexpression lines (Figure 2; (Gazzarrini et al., 2004). This suggests that *bpc1*/*2* phenotypes may be caused by ectopic expression of *FUS3*. To address the genetic relationship between class I *BPCs* and *FUS3*, we crossed *bpc1*/*2* with *fus3*-*3*. The *bpc1*/*2 fus3*-*3* triple mutant indeed showed partial rescue of these phenotypes, including plant height (Figure 6A,D), silique and seed abortion (Figure 6B,C,E,F), as well as embryo sac development (Figure 6H), supporting the hypothesis that *FUS3* is misexpressed in *bpc1/2*.

**Figure 6.**
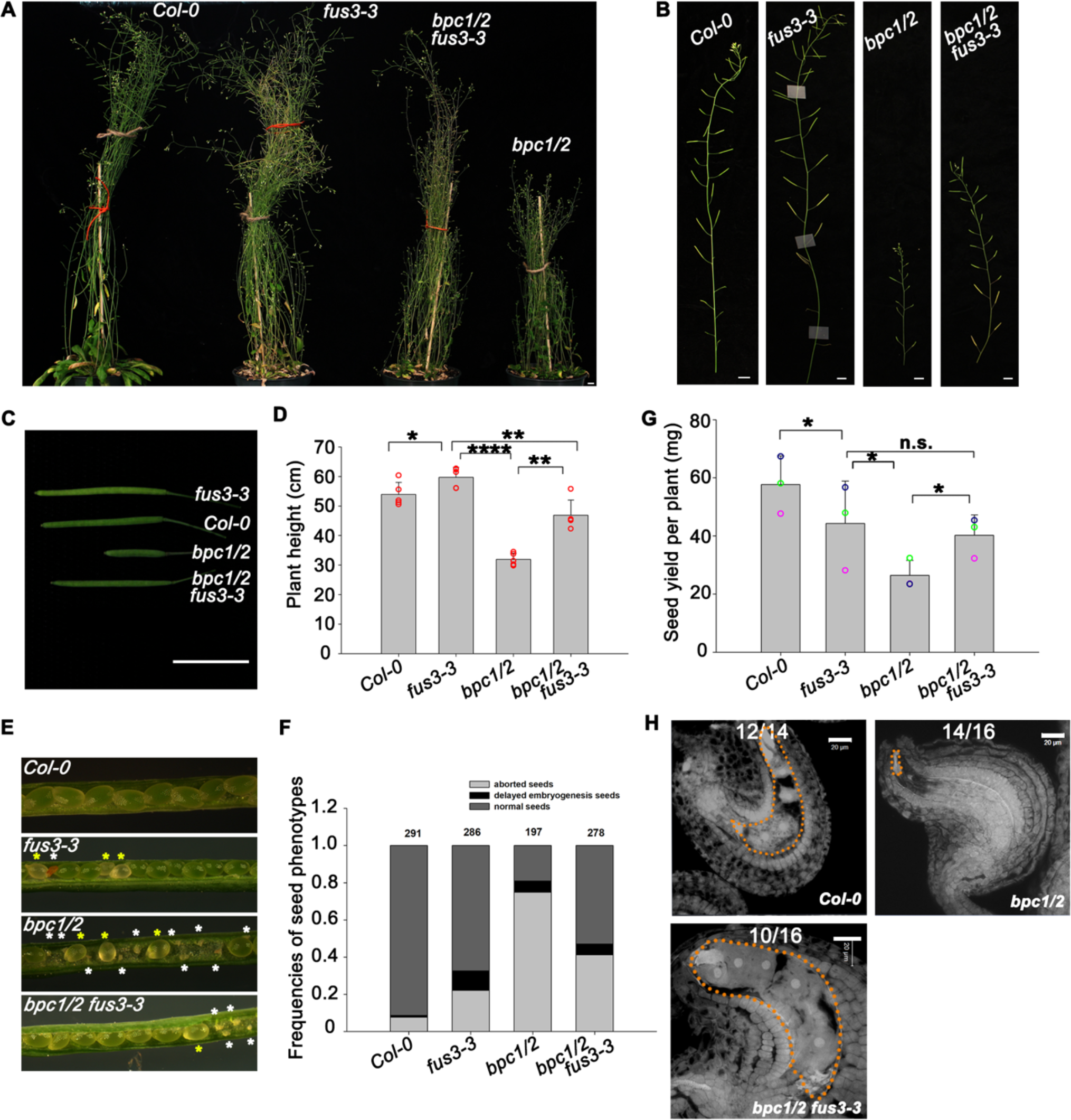
*P*artial rescue of *bpc1*/*2* stunted growth, aborted ovules and seeds in *fus3*-*3 bpc1*/*2*. **A**, The stunted growth of *bpc1*/*2* was partially rescued in *bpc1*/*2 fus3*-*3*. **B, C**, *bpc1*/*2 fus3*-*3* partially rescues *bpc1*/*2* reduced silique elongation. Scale bar, 1cm. **D**, Quantification of the plant height. Five biological replicates were performed. Each replicate consisted of five plants per genotype. **E, F**, *fus3-3* partially rescues *bpc1*/*2* severe seed abortion. The white asterisk in **E** represents aborted seed, while the yellow asterisk represents the delayed embryogenesis seeds. **F**, Frequencies of seed phenotypes in *bpc1*/*2 fus3*-*3* mutants. The total number of sees was calculated in 10 peeled siliques (half side). Three biological repeats were performed with similar results and one is shown (see also Supplemental Figure 10A). **G**, The seed yield of *bpc* mutants. Error bars represent the SD of three biological replicates (n=5). n.s.: no significant difference. (* p<0.05; ** p<0.01; **** p<0.0001); student t-test was used. **H**, *fus3*-*3* partially rescues the embryo sac defects of *bpc1/2*. The image was taken at 1DAP. Scale bar, 20μm. Numbers refer to the number of embryos displaying the phenotype shown.

After fertilization, the endosperm of some *bpc1*/*2* mutants appeared very dense and some ovules were not fertilized (Figure 6H; Supplemental Figure 8E). In fertilized seeds, most *bpc1/2* also display delayed or arrested embryo development (Figure 6E,F,H; Supplemental Figure 8A,B,F,G). Overall, reproductive defects in higher order *bpc* mutants result in severe reduction of seed yield (Figure 6G). The *bpc1*/*2 fus3*-*3* triple mutant partially rescue endosperm and embryo development (Figure 6E,F,H). Thus, these data strongly suggest that BPCs repress *FUS3* during reproductive and seed development.

### BPC1/2 repress *FUS3* to promote inflorescence stem elongation, ovule and endosperm development

To confirm a repressive role of BPCs on *FUS3* function, we analyzed *FUS3* expression level and patterns in *bpc1*/*2* mutants. We show that *FUS3* transcript level is indeed increased in *bpc1*/*2* inflorescence stem (Figure 7A). Consistent with the transcript analysis, *pFUS3:GUS* activity is also increased in *bpc1*/*2* inflorescence stem and flower buds (Figure 7B). In WT, low *FUS3* expression in the inflorescence stem is shown by transcriptomic data and detected with the *pFUS3:FUS3ΔC-GFP* sensitive reporter (Supplemental Figure 1). Together with previous findings showing that plant height is reduced in *pML1:FUS3*-*GFP* misexpression plants (Gazzarrini et al., 2004), while increased in the *fus3*-*3* mutant (Figure 6D), these results indicate that BPC1/2 downregulates *FUS3* in the stem to promote stem elongation.

**Figure 7.**
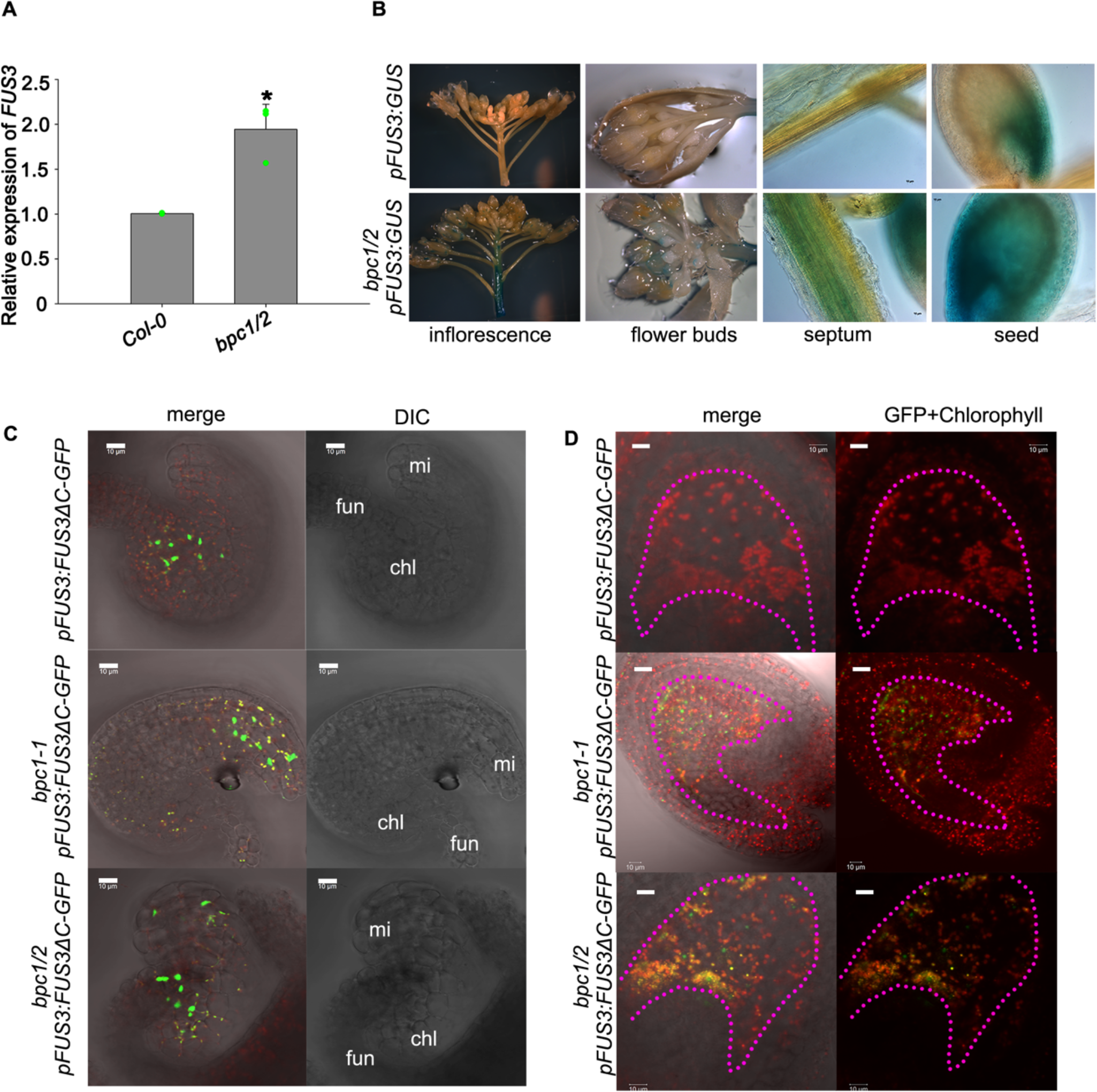
BPC1/2 negatively regulate *FUS3* expression in reproductive organs and seeds. **A**, qRT-PCR showing increased *FUS3* transcript level in *bpc1/2* inflorescence stem. Error bars represent the SD of three biological replicates (* p<0.05; student t-test). **B**, GUS staining in the inflorescence stem, flower buds, septum and seed (2DAF) of *pFUS3:GUS* and *bpc1*/*2 pFUS3:GUS* lines. The GUS staining was enhanced in the inflorescence stem and septum, while ectopically expressed in the endosperm of *bpc1*/*2*. **C, D** *pFUS3:FUS3ΔC*-*GFP* and *bpc1 pFUS3:FUS3ΔC*-*GFP* ovules were images before (C) and two days after (D) fertilization by confocal microscopy. FUS3ΔC-GFP was localized to the chalaza region of developing WT ovules before fertilization, while ectopically localized to the integuments at the micropilar region of *bpc1*-*1 and* of *bpc1*/*2* ovules (FS12) and to the endosperm of 2DAF *bpc1*-*1* seeds.

During reproductive development FUS3ΔC-GFP is mislocalized to the integuments at the micropilar region of developing *bpc1*-*1 and bpc1*/*2* ovules, while after fertilization ectopic *pFUS3:GUS* activity and FUS3ΔC-GFP localization were detected in *bpc1* and and *bpc1*/*2* endosperms (Figure 7B,C,D). Combined with the above functional analysis, these results show that before fertilization BPCs restrict *FUS3* expression to the funiculus and chalazal region of the ovule to promote ovule development, while after fertilization *FUS3* is repressed by BPCs in most of the endosperm to coordinate embryo and endosperm growth.

To analyze the repressive role of class I BPCs, we also crossed *pFUS3:FUS3:GFP* translational reporter with *bpc1*/*2* mutant. However, we were only able to isolate *bpc1*-*1 pFUS3:FUS3:GFP* lines. As shown in Supplemental Figure 8 and previous research (Monfared et al., 2011) *bpc1*-*1* doesn’t have any visible phenotype compared with wild type, nor does *pFUS3:FUS3:GFP*, which rescue the *fus3*-*3* mutant phenotypes (Gazzarrini et al., 2004; Chan et al., 2017). However, in *bpc1*-*1 pFUS3:FUS3:GFP* some flower buds were arrested and never opened, resembling *bpc1*/*2* mutant (Figure 8A). In those flower buds, petal and anther filament did not elongate and anthers were aborted, similar to *bpc1*/*2* double mutant (Figure 8A). Seed abortion was increased and delayed embryogenesis was evident in *bpc1*-*1 pFUS3:FUS3:GFP* plants (Figure 8B,C). The *bpc1*-*1 pFUS3:FUS3:GFP* plants were shorter compared with *bpc1*-*1*, *pFUS3:FUS3:GFP* or wild type and resembled the *bpc1*/*2* double mutant (Figure 8D and Figure 6A,D). Thus, our inability to isolate *bpc1*/*2 pFUS3:FUS3:GFP* mutant may be due to the severe phenotype of such a mutant. The presence of the *pFUS3:FUS3:GFP* transgene enhanced the *bpc1*-*1* phenotype likely due to higher or ectopic *FUS3* expression. Accordingly, we could detect strong GFP fluorescence in the integuments, seed coat and funiculus of *bpc1*-*1 pFUS3:FUS3:GFP*, while *pFUS3:FUS3:GFP* showed no fluorescence in WT (in contrast to the stable *pFUS3:FUS3dC*-*GFP*). Furthermore, FUS3:GFP was mis-localizated in *bpc1*-*1 pFUS3:FUS3:GFP* endosperm after fertilization (Figure 8E), in agreement with *pFUS3:FUS3ΔC-GFP* and *pFUS3:GUS* mislocalization in *bpc1*-*1* and *bpc1/2* endosperm (Figure 7). These results further support a repressive role of BPCs on *FUS3* expression in different tissues during reproductive and seed development.

**Figure 8.**
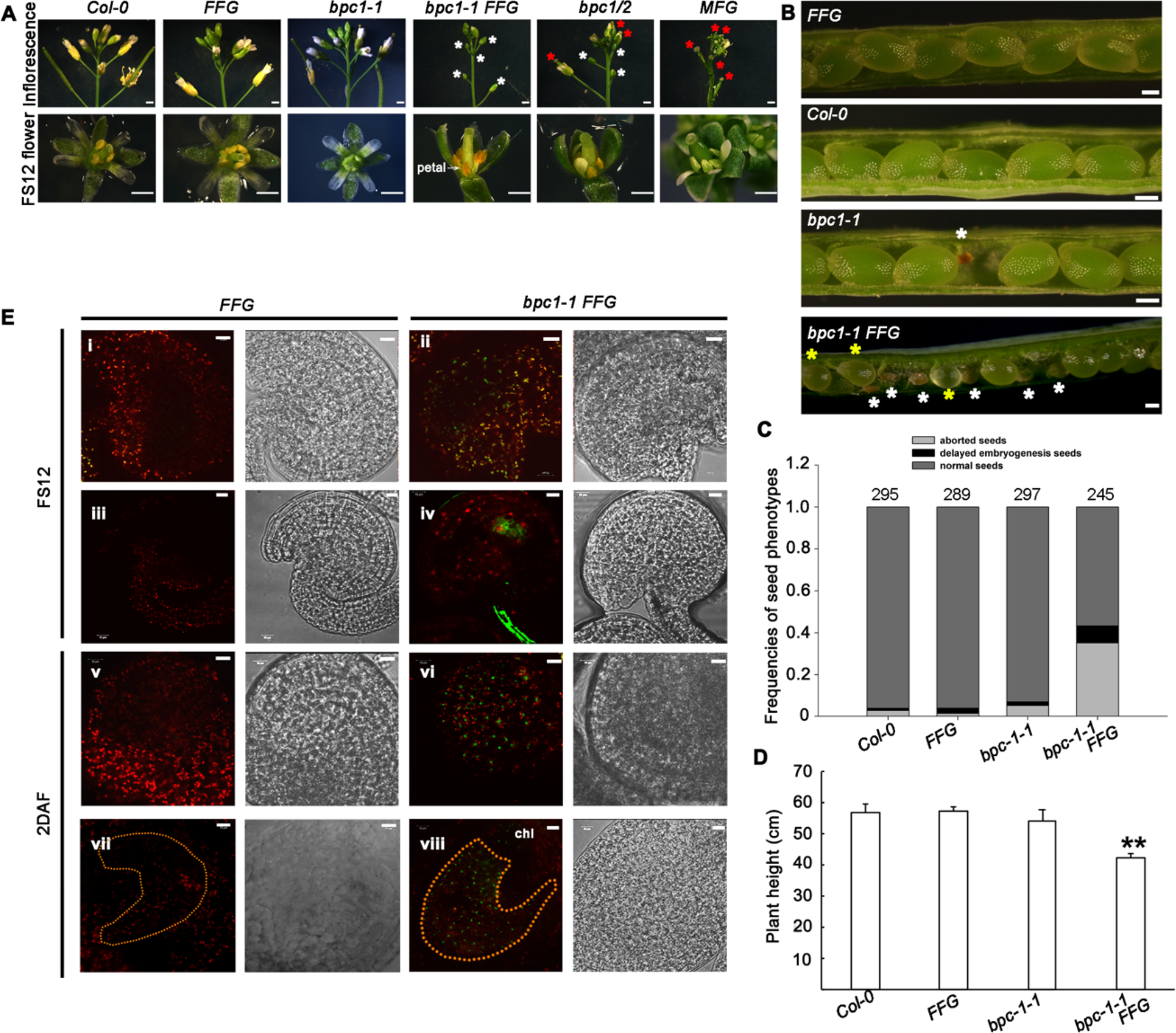
Ectopic *FUS3* expression negatively impacts reproductive organ development. **A**, Introduction of a *pFUS3:FUS3*-*GFP (FFG)* transgene in *bpc1*-*1* mutant results in arrested flower buds that never open (white asterisk), similar to *bpc1*/*2* double mutant. The arrested flower buds in *bpc1*-*1 FFG* have underdeveloped petals, non-elongated filaments and aborted anthers, similar to *bpc1*/*2*. *pML1:FUS3*-*GFP* (*MFE*) also show shorter filaments and underdeveloped anthers, but flower buds open prematurely. **B**, *bpc1*-*1 FFG* mutant caused aborted seeds and delayed embryogenesis. Aborted seeds (white asterisk) and delayed embryogenesis (yellow asterisk) are shown. **C**, Frequencies of seed phenotypes. The total number of seeds was calculated in ten siliques (half side). Three biological repeats were performed, and one representative is shown (see also Supplemental Figure 10B). **D**, *bpc1*-*1 FFG* plants display stunted growth. The error bar represents SD of three biological replicates (n=5). (**: p<0.01; student t-test was used). **E**, *FUS3* is mis-expressed in the integument (**ii**) and increased in the funiculus (**iv**) of *bpc1*-*1* at FS12. Two days after fertilization (2DAF), FUS3-GFP is increased in the seed coat (**vi**) and mis-expressed in the endosperm (**viii**) at 2 DAF.

Upon closer inspection of *bpc1*/*2*, *bpc1*-*1 pFUS3:FUS3:GFP* and *pML1:FUS3*-*GFP* ovules that were successfully fertilized we noticed that they had an increased number of endosperm nuclei, which correlated with an increase in seed size (Figure 9A,B,C,D; Supplemental Figure 9). In fertilized ovules, some embryos were delayed or arrested at various stages (globular to early torpedo) of development compared to wild type (Figure 9E; Supplemental Figure 9). Lastly, *bpc1*/*2* mutants also showed aberrant cell division patterns in the embryo and suspensor, which resulted in defective embryos and were partially rescued by *fus3*-*3* (Figure 9E; Supplemental Figure 9). Collectively, these data show that repression of *FUS3* in the endosperm of developing seeds is required to coordinate endosperm and embryo growth.

**Figure 9.**
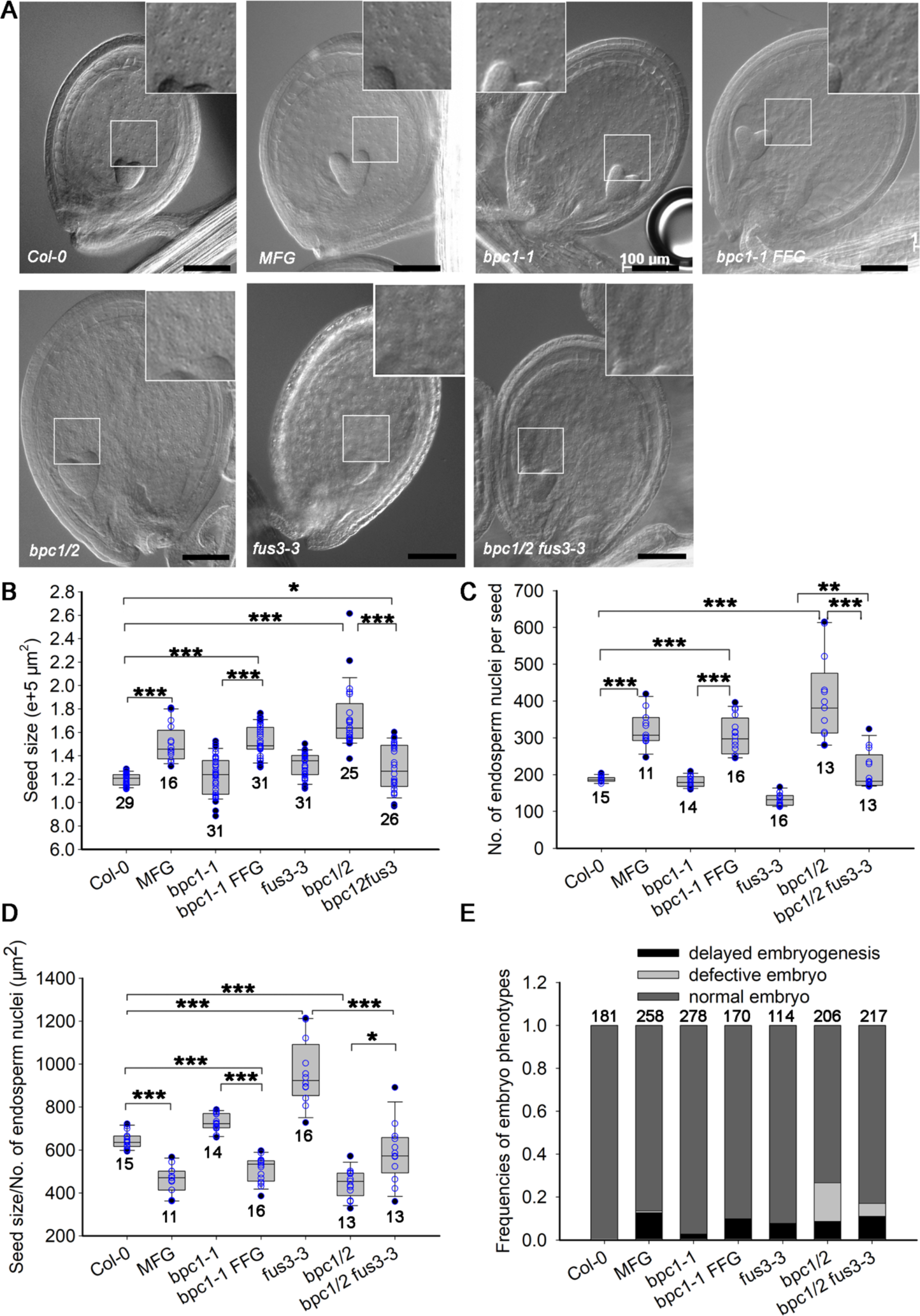
BPC1/2 negatively regulate endosperm nuclei proliferation and seed size by repressing *FUS3*. **A**, Whole-mount clearing, **B**, seed size, **C**, quantification of endosperm nuclei, **D** seed size versus number of endosperm nuclei and **E**, frequencies of embryo phenotypes of wild-type (*Col*-*0*), *pML1:FUS3*-*GFP (MFG)*, *bpc1*-*1*, *bpc1*-*1FFG*, *bpc1*/*2*, *fus3*-*3* and *bpc1*/*2 fus3*-*3* seeds at 6DAP. Over-proliferation of endosperm nuclei and larger seed size in the *bpc1*/*2*, *bpc1*-*1FFG* and *MFG* lines, and partial rescue in *bpc1/2 fus3-3*. Images were taken 6DAP. Scale bar, 100μm. **B**, **C** Ectopic expression of *FUS3* in *MFG*, *bpc1*-*1 FFG*, and *bpc1*/*2* leads to enlarged seed size **B)**, increased endosperm nuclei proliferation **C)** and density **D)**, which is partially rescued in *bpc1*/*2 fus3*-*3*. **E**, Ectopic expression of *FUS3* in *bpc1*/*2*, *bpc1*-*1FFG* and *MFG* results in delayed embryogenesis; *bpc1*/*2* defective embryos are partially rescued by *fus3*-*3*. (* p<0.05; ** p<0.01; **** p<0.0001, student t-test was used).

## DISCUSSION

PRC2 play important roles in balancing cell proliferation with differentiation and regulating developmental phase transitions in plants and animals. Recently, genome wide studies have shown that the plant-specific, class I BPC transcription factors bind Polycomb response elements (PREs), recruit EMF-PRC2 and trigger gene silencing during germination (Xiao et al., 2017). Similar to GAGA factors in *Drosophila melanogaster*, BPCs recognize (GA/CT)_n_ cis elements, despite the lack of sequence similarity between these transcription factors, suggesting convergent evolution (Berger and Dubreucq, 2012). BPCs play essential roles during vegetative and reproductive development, as shown by the dwarf stature and severe seed abortion displayed by higher order *bpc* mutants, however the molecular mechanisms are largely unknown (Kooiker et al., 2005; Monfared et al., 2011; Simonini et al., 2012; Simonini and Kater, 2014;). Here we show that BPC1/2 interact with FIS-PRC2 and bind to the *FUS3* chromatin to restrict *FUS3* expression to specific tissues during reproductive and seed development. BPC-mediated spatiotemporal regulation of *FUS3* expression is required to i) suppress stem elongation during vegetative-to-reproductive phase change, ii) promote ovule development before fertilization and iii) coordinate embryo and endosperm development after fertilization (Figure 10). Several lines of evidence support these conclusions. First, Y1H show that class I BPCs bind to (GA/CT)_n_ repeats around the *FUS3* transcription start, and ChIP assays in flower buds show that BPC1 binds *in vivo* to the *FUS3* chromatin. Mutations in these (GA/CT)_n_ sites abolish BPCs binding and derepress *FUS3* during vegetative development. Furthermore, *FUS3* is upregulated in the inflorescence stem of *bpc1*/*2* dwarf plants, which is consistent with *fus3*-*3* tall plant and *ML1:FUS3*-*GFP* dwarf plant phenotypes, as well as FUS3 role as repressor of vegetative-to-reproductive phase change (Gazzarrini et al., 2004; Lumba et al., 2012). Second, class I BPCs interact with FIS2-PRC2 complex *in planta*, and the *in vivo* BPC1-binding region on *FUS3* was shown to associate with MEA and H3K27me3 repressive marks (Makarevich et al., 2006), strongly suggesting BPC1 recruits FIS-PRC2 to repress *FUS3* during reproductive/seed development. Third, FUS3 is transiently localized to the integuments during early ovule development and later restricted to the funiculus and chalaza of mature wild type ovules. Ectopic and persistent expression of *FUS3* in the integuments of *bpc1*/*2* and *ML1:FUS3* mis-expression lines impairs integument and embryo sac development leading to seed abortion, which can be partially rescued in *fus3-3 bpc1*/*2*. Last, after fertilization FUS3 is localized to the funiculus, chalaza and outer integument, aside from its known localization to the embryo (Gazzarrini et al., 2004). Ectopic expression of *FUS3* in *bpc1*/*2* and *ML1:FUS3* endosperm leads to increased proliferation of the endosperm nuclei and delayed or arrested embryo development, which are rescued in *fus3*-*3 bpc1*/*2*. The latter phenotypes are also displayed by mutants in *FIS-PRC2* subunits (Kiyosue et al., 1999; Kohler and Grossniklaus, 2002). We conclude that BPCs recruit PRC2 to restrict spatiotemporal *FUS3* expression during reproductive and seed development; this is required to regulate tissue development locally and modulate developmental phase transitions in Arabidopsis. The genomic sequences of *FUS3* orthologs in other species show conservation of (GA/CT)_n_ repeats (Supplemental Figure 11), suggesting that similar mechanisms may regulate the expression of *FUS3*-like transcription factors in other species.

**Figure 10.**
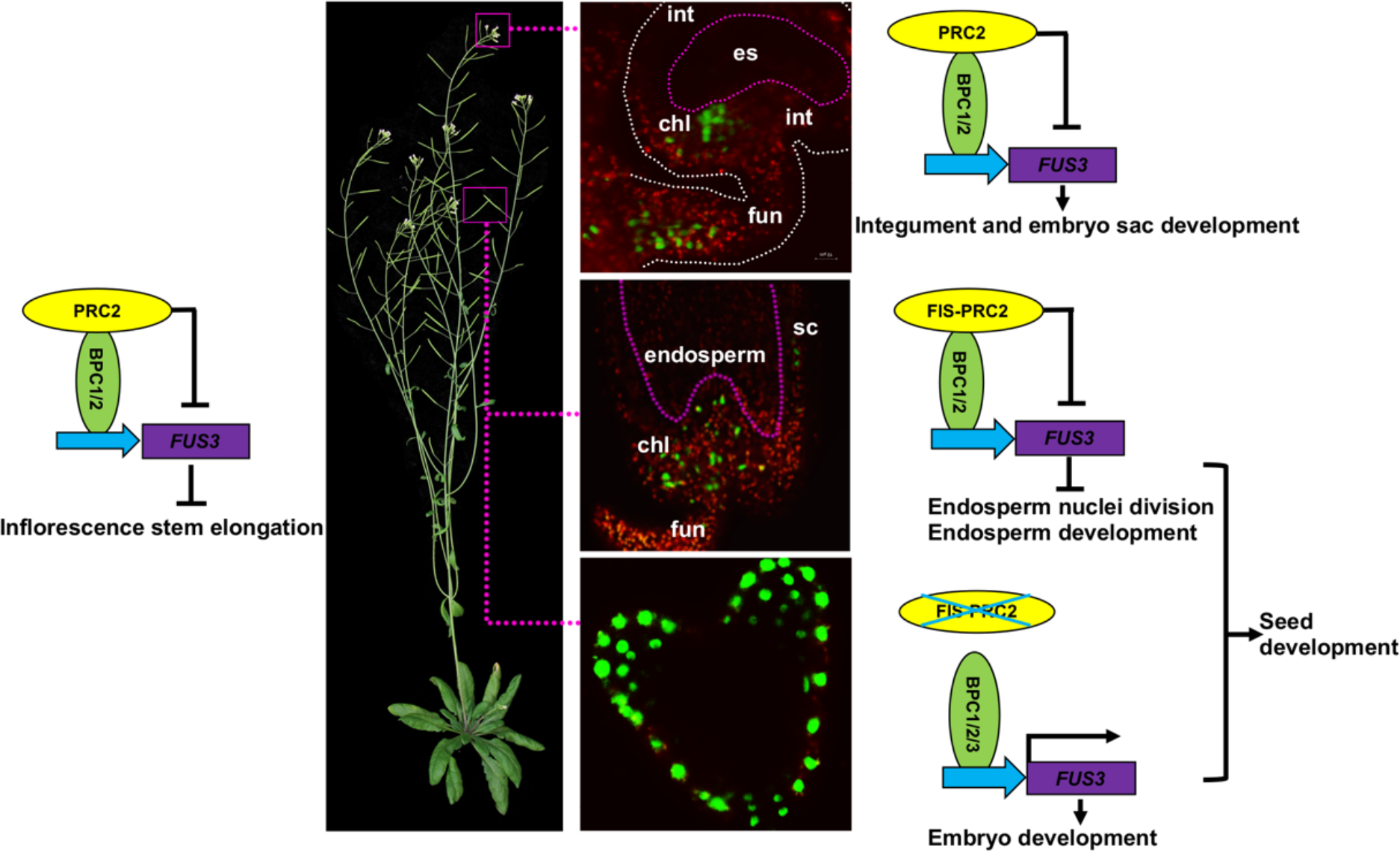
Spatiotemporal restriction of *FUS3* expression by BPC1/2 during reproductive and seed development. Model depicting spatiotemporal expression of *FUS3* and its role in the regulation of vegetative-to-reproductive and gametophytic-to-sporophytic phase transitions. During the vegetative-to-reproductive phase change, *FUS3* is repressed by BPC1/2 in the inflorescence stem to allow stem elongation. During ovule development, FUS3 becomes restricted to the funiculus and chalaza through BPC1/2-mediated repression in the integuments; this is required to promote integument and embryo sac development. After fertilization, FUS3 is localized to the embryo, seed coat, chalaza and funiculus, but is repressed in the endosperm by BPC1/2 to decrease endosperm nuclei division and promote embryo development. In the stem, BPC1/2-mediated *FUS3* repression may be orchestrated by EMF-PRC2, which interacts with BPC1/2 and represses *FUS3* postembryonically (Liu et al., 2016; Xiao et al., 2017). FUS3 repression in the integuments may require sporophytic VRN/EMF PRC2. After fertilization, FIS-PRC2 represses *FUS3* in the endosperm (Makarevich et al., 2006).

### Inflorescence stem elongation and flower development require repression of *FUS3* by Class I BPCs

During germination BPC1 directly binds to the genomic region of *FUS3* proximal to the transcription start, which is marked by H3K27me3 repressive marks and associates with FIE (Figure 3; Xiao et al., 2017). Furthermore, *FUS3* is strongly expressed in *swn clf* seedlings (Makarevich et al., 2006), suggesting that during germination FUS3 is repressed through BPC1-recruitment of EMF/VRN-PRC2. Here we show that mutations of all BPC binding sites on the *FUS3* promoter derepress *FUS3* in vegetative tissues, and that lack of BPCs results in ectopic *FUS3* expression in leaves, inflorescence stem and flower buds. Furthermore, ectopic *FUS3* in *bpc1*/*2*, *bpc1 pFUS3:FUS3*-*GFP* or *pML1:FUS3*-*GFP* leads to similar phenotypes, including reduced internode elongation and defective flowers (arrested flower bud development, flowers with a protruding carpel and shorter floral organs), suggesting *FUS3* inhibits the elongation of the stem and floral organs during flowering. Recently, deletion of a small region in the *FUS3* promoter near the BPC binding sites and corresponding to the PRC2 recruitment region, lead to ectopic *FUS3* expression in vegetative tissues (Roscoe et al., 2019). Thus, we propose that class I BPCs recruit VRN/EMF-PRC2 to repress *FUS3* post-embryonically, more specifically in germinating seeds, in vegetative and reproductive organs (Figure 10).

Although *bpc1*/*2* shows dramatic phenotypes during reproductive development, germination and early seedling development are not affected as it would be expected from derepression of embryonic genes. This may be due to functional redundancy within the BPCs family and the difficulty in isolating and characterizing higher order *bpc* mutant due to sterility (Monfared et al., 2011). However, the C1-2iD ZnF TF AZF1 associates with *LEC2*, *FUS3* and *ABI3* genomic regions that are also bound by BPC1 and that colocalize with FIE and PRC2 H3K27me3 peaks, suggesting that PRC2-dependent *FUS3* and *LAFL* gene silencing during post-embryonic development requires BPC, ZnF and likely other factors (Xiao et al., 2017; Zhou et al., 2018). This is also consistent with the strong phenotype shown by telobox binding mutants (*trb1*/*2*/*3*), which is enhanced by mutations in PRC2 (Zhou et al., 2018). Given that 42% of genome-wide FIE association regions were bound by BPC1 and AZF1, a combinatorial role for these transcription factors in recruiting PRC2 and triggering gene silencing has been proposed (Xiao et al., 2017; Zhou et al., 2018).

### BPC-mediated restriction of *FUS3* expression in developing ovules and seeds is required to promote ovule development and to coordinate endosperm and embryo growth

During ovule development, the funiculus supplies nutrients and signaling molecules from the mother plant to the chalaza, initiates the integuments that grow around the nucellus and protect the developing female gametophyte (Schneitz et al., 1995). Our data show that during megagametogenesis FUS3 is initially localized to the nucellus epidermis and tissues surrounding the nucellus, including the integuments and chalaza. However, BPC1/2 later repress *FUS3* in the integuments of mature ovules, and ectopic *FUS3* expression in *bpc1*/*2* inhibits integuments and embryo sac development, triggering ovule abortion. These phenotypes are recapitulated in *pML1:FUS3* misexpression lines, where the *pML1* promoter specifically drives expression of *FUS3* in the integuments and endothelium also in mature ovules, but rescued in *bpc1*/*2 fus3*-*3* mutant, strongly indicating that spatiotemporal restriction of FUS3 localization is required for integuments, embryo sac and ovule development (Figure 10). This is in agreement with previous finding showing that the integuments are required for female gametogenesis (Elliott et al., 1996; Klucher et al., 1996; Baker et al., 1997).

Following fertilization, the zygote together with the endosperm and the integuments develop in a coordinated manner to form the embryo and the seed coat of the mature seed. FUS3 was previously shown to localize to developing embryos from globular to cotyledon stages (Gazzarrini et al., 2004). Using the sensitive/stable FUS3dC-GFP reporter, we found that FUS3 localizes also to the funiculus, chalaza and outer seed coat of developing seeds, partially mirroring its expression pattern in ovules. In *bpc1*/*2* mutant or in *pML1:FUS3*-*GFP* misexpression lines, ectopic FUS3 localization to the endosperm increases cell proliferation resulting in enlarged endosperm and larger seeds at the expense of embryo development, which is typically delayed or arrested in *bpc1*/*2* and *pML1:FUS3*-*GFP* compared to WT. These phenotypes are reminiscent of some FIS-PRC2 mutant alleles of *mea* (Kiyosue et al., 1999). Given that *FUS3* is derepressed in *mea* endosperm and that MEA and H3K27me3 repressive marks associate in a repressive region of the *FUS3* locus where BPC1 also binds, we propose that BPC1/2 recruit FIS-PRC2 to repress *FUS3* in the endosperm (Makarevich et al., 2006); this is required to reduce the rate of endosperm nuclei proliferation, promoting endosperm differentiation and embryo growth (Figure 10).

In the absence of fertilization seed development is repressed by PRC2. FIS-PRC2 represses autonomous central cell division in the ovule and regulate endosperm development after fertilization. The FIS-PRC2 specific subunits, MEA and FIS2, are targeted solely to the central cell in the ovule and endosperm in the seed, and thus are likely to participate in *FUS3* repression in these tissues (Luo et al., 2000; Wang et al., 2006). The MEA homolog SWN, which belongs to the VRN-PRC2 and FIS-PRC2 complexes, has a broader localization pattern, but plays a partially redundant function with MEA in repressing central cell/endosperm nuclei proliferation in the absence of fertilization (Wang et al., 2006). Thus, SWN may also be involved in repressing *FUS3* in the central cell/endosperm. In contrast, autonomous seed coat development in the ovule is repressed by the sporophytic complexes VRN-PRC2 and EMF-PRC2, which may be involved in repressing *FUS3* in the integuments (Kohler and Grossniklaus, 2002; Roszak and Kohler, 2011). In accordance, *FUS3* and other seed-specific genes were derepressed and showed reduced H3K27me3 repressive marks in siliques of a weak *curly leaf* (*clf)* allele, although the tissue specific expression was not investigated (Liu et al., 2016).

Although BPCs can recruit EMF- and FIS-PRC2 complexes for transcriptional silencing, BPCs were also shown to positively regulate a close *FUS3* family member, *LEC2* (Berger et al., 2011). This is in accordance with the role of GAGA binding proteins in animals, which have dual function of activators and repressors (Berger and Dubreucq, 2012). Interestingly, FUS3 is expressed in the embryo and in specific sporophytic tissues of the ovule and seed (chalaza, funiculus, seed coat), where all class I BPCs are expressed. Thus, it will be important to determine the mechanisms of BPCs activation and repression of *FUS3* and other *LAFL* genes during reproductive and seed development.

Collectively, these findings indicate that spatiotemporal restriction of *FUS3* expression is necessary for organ development and to allow the transition between various phases of development. An important question is how does FUS3 regulate tissue development and phase transitions. FUS3 was shown to be a nexus in hormone synthesis; by controlling the ABA/GA ratio, FUS3 promotes seed maturation while inhibiting germination and flowering, with ABA and GA acting as positive and negative regulators of FUS3 protein levels, respectively (Gazzarrini et al., 2004; Lu et al., 2010; Chiu et al., 2012). A positive feedback regulatory loop has been established also between auxin and FUS3 in the embryo, whereby FUS3 promotes auxin synthesis and auxin induces *FUS3* (Gazzarrini et al., 2004). Several studies have shown that *LAFL* genes are involved in regulating auxin biosynthesis, which also ties to their role in somatic embryogenesis (Lepiniec et al., 2018). Given that auxin is required for the synchronized growth of the fruit, the different tissues within the seed (integuments, endosperm and embryo) and that FUS3 localization patterns in ovules and seeds largely mirror those of auxin, we propose that FUS3 may regulate auxin level/localization and that auxin may in turn regulate FUS3 expression/activity (Gazzarrini et al., 2004; Figueiredo et al., 2015; Figueiredo et al., 2016; Larsson et al., 2017; Robert et al., 2018). Reduced auxin accumulation in the chalaza and funiculus of *fus3*-*3* or increased auxin levels in the integuments and endosperm of *pML1:FUS3* or *bpc1*/*2* would impair ovule and seed development resulting in seed abortion and delayed embryo development, respectively, as shown by delayed endosperm cellularization and embryo growth arrest triggered by auxin overproduction in the endosperm (Figueiredo and Kohler, 2018; Batista et al., 2019; Robert, 2019).

In conclusion, mutations affecting FIS-PRC2 or PRE binding TF BPCs cause severe seed abortion, however the molecular mechanisms are still poorly understood (Monfared et al., 2011; Wang and Kohler, 2017; Figueiredo and Kohler, 2018). Here we show that BPC1/2-mediated spatiotemporal restriction of *FUS3*, a target of the PRC2 complex, is required for the development of ovule and seed tissues and to regulate developmental phase transitions.

## MATERIAL AND METHODS

### Plant material

T-DNA insertion lines *bpc1*-*1* (SALK_072966C), *bpc2* (SALK_090810), *bpc1*-*1*/*bpc2* (*bpc1*/*2*; CS68700), and *bpc1*-*1*/*bpc2*/*bpc3*-*1* (CS68699), and an EMS mutant *bpc3*-*1* (CS68805) were previously described (Monfared et al., 2011). T-DNA insertion lines *bpc1*_*salk* (SALK_101466C), *bpc2*_*salk* (SALK_110830C), *bpc3_sail* (SAILseq_553_B09.0) were obtained from ABRC. All primers used for genotyping are listed in the Supplemental Table 1. The *pFIE:FIE:GFP*, *pMSI1:MSI1:GFP*, *pMEA:MEA:YFP* and *pFIS2:GUS* reporter lines were previously described (de Lucas et al., 2016). The *pML1:FUS3-GFP* construct previously described (Gazzarrini et al., 2004) was transform into *fus3*-*3* loss-of-function mutant (Keith et al., 1994). *pFUS3:FUS3ΔC-GFP* construct previously described was transformed into *Col*-*0* (Lu et al., 2010). *pFUS3:FUS3*-*GFP* construct was previously described (Gazzarrini et al., 2004). For transgenic plants carrying the (GA/CT)_n_ mutant promoter reporter [*pFUS3(1.5kb*), 1.5kb upstream of *FUS3* coding sequence with or without mutated (GA/CT)_n_ motifs (shown in Supplemental Figure 2)] was PCR amplified (primers listed in Supplemental Table 1) and cloned into pCAMBIA1391-GUS and pCAMBIA1391-GFP vectors by restriction enzyme digestion (*Hind* III and *BamH* I). Eight to ten transgenic lines per constructs were selected on MS containing 30mg/L hygromycin plates and analyzed for GUS staining or GFP fluorescence. Sterilized Arabidopsis seeds were germinated on half-strength Murashige and Skoog (MS) medium, transferred to soil and grown under 16/8h light/darkness 22°C/18°C. Frequencies of seed phenotypes displayed by various genotypes were calculated with half dissected siliques (n=10); experiments were repeated three times with similar results and one is shown. Total seed yield per plant was calculated with 5 plants per pot, experiments were repeated three times.

### Yeast one-hybrid screening

Yeast one-hybrid library screening and one-on-one retests were performed as described by Deplancke et al. (2006), with some modifications. To construct the baits, 615bp of the *FUS3* genomic sequence upstream of the translation start [*pFUS3(0.6kb);* base pairs −615 to +1), or the *pFUS3* with the mutated (GA/CT)_n_ motif [*pFUS3^MUT^(0.6kb)*] or the truncated *pFUS3* (F1 to F4) were PCR-amplified and recombined into the pDEST-HISi-2 vector by Gateway cloning. Mutagenesis of (GA/CT)_n_ motifs on *pFUS3*^*MUT*^*(0.6kb)* was generated by PCR-driven overlap extension (Heckman and Pease, 2007) with primers listed in Supplemental Table 1. The linearized vectors (digested by *XhoI*) were then transformed into the yeast strain YM4271(a) using the LiAc/SS carrier DNA/PEG method (Gietz and Schiestl, 2007). Transformed yeast colonies were tested for background expression of the *HIS3* reporter and the appropriate 3-aminotriazole (3-AT) concentration was selected. An *Arabidopsis thaliana* transcription factor library (Mitsuda et al., 2010) was transformed into the yeast strain EGY48(α) by electroporation. The initial screening was performed by mating YM4271(a) containing the bait *pFUS3* (0.6kb) with EGY48(α) containing the library on YPD plates overnight. Colonies were selected on medium without Ura, His and Leu, supplemented with 20mM 3-AT (SDA-Ura-His-Leu + 20mM 3-AT). Plasmids isolated from 69 out of 200,000 CFU harbored *BPC3*. To test the binding preference of BPC1-3 on the *FUS3* promoter, BPCs were PCR-amplified and recombined into pDEST-GAD424 by Gateway. The recombined vectors were then transformed into yeast strain EGY48(α). A single transformed YM4271(a) colony containing different truncated or mutated promoters [F1 to F4 and *pFUS3*^*MUT*^*(0.6kb)*, described above] was used for mating with EGY48(α) containing BPCs. Mating and selection procedures were described in Wu et al. (2018). The interaction was judged by the growth of yeast on selection media on the third day.

### Bimolecular fluorescence complementation (BiFC) assay

The CDS of *BPCs*, *FIE*, *MSI1*, *MEA* and *FIS2* were cloned into BiFC vectors pB7WGYN2 (YNE) or pB7WGYC2 (YCE) (Tsuda et al., 2017) by Gateway. These recombined vectors were transformed into *Agrobacterium tumefaciens* strain GV2260 and infiltrated into *Nicotiana benthamiana* leaves as described previously (Duong et al., 2017). At least three biological replicates were performed.

### Differential interference contrast (DIC) microscopy

Pistils at FS12 or siliques were dissected and immersed in fixing solution (9:1, ethanol:acetic acid, v/v) for 2h before washing them twice with 90% ethanol. The siliques were then cleared with clearing solution (2.5g/ml chloral hydrate and 30% glycerol) overnight. Images were taken by a Zess Axioplant 2 microscope equipped with DIC optics. The quantification of seed size and endosperm nuclei are performed by Image J software.

### Confocal microcopy

To observe the expression of GFP signal in transgenic Arabidopsis, fresh tissues was dissected and mounted on the slides with 10% glycerol. Visualization was done with a Zeiss LSM510 confocal microscope (488 nm excitation and a 515-535 nm band pass filter).

### GUS staining

The *pBPC3:GUS* line was previously described (Monfared et al., 2011). The promoter regions of *BPC1*/*2* described in Monfared et al. (2011) were PCR-amplified and transformed into the pGWB3 vector to generate *pBPC1:GUS* and *pBPC2:GUS*. Several transformed homozygous lines were selected on kanamycin and hygromycin plates and analyzed and two lines were selected for further analysis. The GUS staining assays were performed as previously described (Wu et al., 2019) with some modifications. The concentration of ferri/ferrocyanide used for *pBPC3:GUS* was 2mM, while 5mM was used for *pBPC1:GUS* and *pBPC2:GUS*. To detect low expression of *FUS3* in inflorescences, leaves or flowers of *pFUS3(1.5 kb):GUS* and *pFUS3*^*MUT*^*(1.5kb):GUS* lines, ferri/ferrocyanide was not included in the buffer. Cleared tissues were imaged by DIC microscopy using Zeiss Axioplant 2.

### Glutaraldehyde staining

To visualize ovule/seed structures, whole pistils/siliques at FS12 or 1-2DAF were fixed in 3% paraformaldehyde in PBS for 15min at room temperature and rinsed twice with PBS. The treated tissues were stained in 5% glutaraldehyde in PBS at 4°C overnight in the dark. Tissues were washed 3 times with PBS and cleared for about 1 to 2 weeks with ClearSee buffer (Kurihara et al., 2015). The images were photographed with a Zeiss LSM510 confocal microscope (530nm excitation and a 560nm long pass filter).

### Gene expression assay

RNA was extracted using the RNeasy Plant Mini Kit (Qiagen). About 1µg of RNA was used for reverse transcription. Quantitive real-time PCR was performed using Step One Plus real-time PCR system (Applied Biosystems) with SYBR premix. *PP2AA3* was chosen as the internal reference gene. Primers used are listed in Supplemental Table 1. Three biological replicates were performed.

### ChIP assay

To generate *35S:BPC1*-*RFP*, the *BPC1* coding sequence was first cloned into pDONR221 (Life Technologies) and subsequently transferred to pB7RWG2 (Flanders Interuniversity Institute for Biotechnology, Gent, Belgium). Arabidopsis plants were transformed with the *35S:BPC1*-*RFP* using the Agrobacterium tumefaciens-mediated floral dip method (Clough and Bent, 1998). Transformant plants were sown on soil and selected by BASTA; the presence of the construct was assessed by genotyping and analysis of RFP expression. Arabidopsis plants were directly sown on soil and kept under short-day conditions for 2 weeks (22°C, 8h light and 16h dark) and then moved to long-day conditions (22°C, 16h light and 8h dark). ChIP assays were performed as described by Gregis et al. (2009) using for BPC1-RFP an anti-RFP V_H_H coupled to magnetic agarose beads RFP-trap_MA® (Chromotek). Real-time PCR assays were performed to determine the enrichment of the fragments. The detection was performed in triplicate using the iQ SYBR Green Supermix (Bio-Rad) and the Bio-Rad iCycler iQ Optical System (software version 3.0a), with the primers listed in Supplemental Table 1. ChIP-qPCR experiments and relative enrichments were calculated as reported by Gregis et al. (2009).

### Accession Numbers

Sequence data from this article can be found in the Arabidopsis Genome Initiative or GenBank/EMBL databases under the following accession numbers: *FUSCA3* (At3g26790), *BPC1* (AT2G01930), *BPC2* (AT1G14685), *BPC3* (AT1G68120), *BPC4* (AT2G21240), *FIE* (AT3G20740), *MSI1* (AT5G58230), *MEA* (AT1G02580), *FIS2* (AT2G35670) and *LHP1* (AT5G17690).

## SUPPLEMENTAL DATA

**Supplemental Figure 1 *FUS3* expression profile in reproductive tissues and stem epidermis.** The images were generated by the eFP browser (www.bar.utoronto.ca) using microarray data (Schmid et al., 2005; Suh et al., 2005; Swanson et al., 2005; Dean et al., 2011). **A**, *FUS3* expression profile in the pistil and different stages of flowers. **B**, The expression profile of *FUS3* in the seed coat at the heart, bending and walking embryo stage. **C**, The expression pattern of *FUS3* in the stem epidermis. **D**, *FUS3* transcripts are higher at the top of the inflorescence stem compared to bottom (*: p<0.05; student t-test). **E**, *pFUS3:FUS3ΔC*-*GFP* fluorescence appears stronger in the epidermis of internodes closer to the flower buds and weaker at the bottom of the stem. Identical confocal settings were used.

**Supplemental Figure 2 Location of (GA/CT)_n_ mutated in the *pFUS3* genomic region used in Y1H.** The (GA/CT)_n_ motifs that were mutated in the *FUS3* sequence upstream of the start codon (base pairs −1 to −615) are highlighted in yellow. Mutated G/C residues are marked in red. Telo boxes binding sites of AZF1 (Xiao et al., 2017) are marked in blue. The repressive region (126bp) that associated with MEA and H3K27me3 repressive marks (Makarevich et al., 2006), and was deleted in promoter studies Roscoe et al. (2019) is underlined.

**Supplemental Figure 3 Class I BPC family members form homo- or hetero- dimers.** BPC1, BPC2 and BPC3 form heterodimers with each other. Only BPC2 and BPC3 form homodimers. Lack of interaction between FUS3 and BPCs or FIS-PRC2 in BiFC assays is shown as the negative control in Supplemental Figure 5.

**Supplemental Figure 4 PRC2 complexes in Arabidopsis.** There are three different types of PRC2 in Arabidopsis depending on subunit composition - VRN-PRC2, EMF-PRC2 and FIS-PRC2-, which regulate vernalization, vegetative development and female gametophyte/seed development, respectively (Mozgova et al., 2015).

**Supplemental Figure 5 FUS3 protein does not interact with FIS-PRC2 complex and Class I BPC family members using BiFC.** Negative control showing that FUS3 does not interact with FIE, MSI1, MEA, FIS2, BPC1, BPC2 or BPC3 in *N. benthamiana* by BiFC.

**Supplemental Figure 6 Class I BPC proteins do not interact with LHP1 using BiFC.** Class I BPC proteins do not interact with LHP1 in *N. benthamiana* by BiFC.

**Supplemental Figure 7 Expression patterns of FIS-PRC2 in ovules and embryos.** FIS2 complex (FIE, MSI1, MEA and FIS2) expression/localization patterns in ovules before fertilization (FG4 and FG7), 1DAP seeds and embyos at globular, heart, torpedo and cotyledon stages. Ant: antipodals; cc: central cell; chl: chalaza; cze: chalazal endosperm; ec: egg cell; fu: funiculus; ii: inner integuments; mce: micropilar endosperm; oi: outer integuments; pen: peripheral endosperm; pge: pre-globular embryo; sdc: seed coat; sus: suspensor; syn: synergids cell. Pink dashed lines represent the outline of the embryo sac. The ferro/ferricyanide used in GUS staining buffer was 2mM for *pFIS2:GUS*.

**Supplemental Figure 8 Class I *bpc* mutants show delayed megagametogenesis, seed abortion and delayed embryogenesis. A**, Class I BPC mutants show seed abortion and delayed embryogenesis phenotypes. The white asterisks indicate aborted seeds; the yellow asterisks represent delayed embryogenesis seeds. **B**, The frequencies of seed phenotypes in 10 peeled half-side of *bpc* mutants siliques. Three biological repeats were performed and one representative result is shown. **C**, Seed yield of WT and *bpc* mutants. The error bars represent the SD of three biological replicates (*: p<0.05; student t-test was used) **D**, FS12 ovules of Class I BPC mutants showing aborted embryo sac and delayed megagametyogenesis. **E**, Class I BPC mutants show condensed endosperm and unfertilized egg cell with the degenerated synergid cell at 2DAF. **F**, Arrested embryos in *bpc1*/*2* with an enlarged seed at 3DAP. **G**, At mature stage (11DAP), some *bpc1*/*2* embryos were arrested at torpedo stage. The pink arrow points to the aborted embryo sac. Ant: antipodals; cc: central cell; cze: chalazal endosperm; ec: egg cell; es: embryo sac; mce: micropilar endosperm; nu: nuclei; pen: peripheral endosperm; dsyn: degenesrated synergid cell; z: zygote. Pink dashed lines outline the embryo sac at FS12; the yellow dash lines outline the embryo sac at 2DAF.

**Supplemental Figure 9 Overexpression of *FUS3* results in embryo defect and over-proliferation of the endosperm nuclei.** Seeds of (**A**) Wildtype, (**B**) *bpc1*-*1*, (**C, D**) *MFG*, (**E, F**) *bpc1*-*1 FFG* and (**G-L**) *bpc1*/*2* at 3 DAP. Pink dashed lines represent the outline of the embryo. The yellow dashed line represents the embryo sac. White arrows indicate the abnormal suspensors. White, yellow or red asterisk indicates the aborted seed, arrested embryo or defective embryo, respectively.

**Supplemental Figure 10 Frequencies of seed phenotypes**. **A, B** The total number of seeds displaying various phenotypes was calculated in 10 peeled siliques (half side) of **A**, WT, *bpc1*/*2* and *bpc1*/*2 fus3* mutants, and **B**, WT, *bpc1*-*1* and *bpc1*-*1 FFG* mutants. Three biological repeats were performed, and two are shown here. See also Figures 6 and 8.

**Supplementary Figure 11 Conserved (GA/CT)_n_ motifs in orthologous** *FUS3* **genes.** Accession number of *FUS3* orthologous genes: *OsLFL1* (LEC2 and FUSCA3-like protein 1;GenBank: EF521182.1) from *Oryza sativa*, *BnFUS3* (NCBI: XM_013792060) from *Brassica napus*, *GmFUS3* (Gene ID: LOC100813055) from *Glycine max*.

**Supplemental Table 1 Primers used in this study.**

## ACKNOWLEDGMENT

We thank C.S. Gasser (UC Davis) for *pBPC3:GUS* reporter; F. Parcy for *pFUS3:GUS*; SM Brady (UC Davis) and M. De Lucas (Durham University) for *pFIE:FIE:GFP*, *pMSI1:MSI1:GFP*, *pMEA:MEA:YFP* and *pFIS2:GUS* reporter lines as well as *FIE* and *MSI1* vectors; C. Koehler (The Swedish University of Agricultural Sciences) and R. Yadegari (University of Arizona) for *MEA*/*pBluescript II KS* and *FIS2*/*pGBKT7* vectors. JW was supported by the National Natural Science Foundation projects (grants 31701952) and China Postdoctoral Council scholarships. V.G. was supported by the Ministero dell’Istruzione, dell’Università e della Ricerca MIUR, SIR2014 MADSMEC, Proposal number RBSI14BTZR. R.P. was supported by the Doctorate School in Molecular and Cellular Biology, Università degli Studi di Milano Fellowship. This work was funded by a Natural Sciences and Engineering Research Council of Canada Discovery Grant to SG.

## AUTHOR CONTRIBUTIONS

JW and SG conceived the study and wrote the paper. JW conducted most of the experiments. SD helped with the identification of higher order mutants. RP and VG conducted ChIP assays. All read and approved the manuscript.

## REFERENCES

Baker SC, Robinson-Beers K, Villanueva JM, Gaiser JC, and Gasser CS. 1997. Interactions among genes regulating ovule development in Arabidopsis thaliana. Genetics 145:1109–1124

Batista RA, Figueiredo DD, Santos-Gonzalez J, and Kohler C. 2019. Auxin regulates endosperm cellularization in Arabidopsis. Genes Dev 33:466–476

Berger N, and Dubreucq B. 2012. Evolution goes GAGA: GAGA binding proteins across kingdoms. Biochim Biophys Acta 1819:863–868

Berger N, Dubreucq B, Roudier F, Dubos C, and Lepiniec L. 2011. Transcriptional regulation of Arabidopsis LEAFY COTYLEDON2 involves RLE, a cis-element that regulates trimethylation of histone H3 at lysine-27. Plant Cell 23:4065–4078

Braybrook SA, Stone SL, Park S, Bui AQ, Le BH, Fischer RL, Goldberg RB, and Harada JJ. 2006. Genes directly regulated by LEAFY COTYLEDON2 provide insight into the control of embryo maturation and somatic embryogenesis. Proc Natl Acad Sci USA 103:3468–3473

Carbonero P, Iglesias-Fernandez R, and Vicente-Carbajosa J. 2017. The AFL subfamily of B3 transcription factors: evolution and function in angiosperm seeds. J Exp Bot 68:871–880

Chan A, Carianopol C, Tsai AYL, Varatharajah K, Chiu RS, and Gazzarrini S. 2017. SnRK1 phosphorylation of FUSCA3 positively regulates embryogenesis, seed yield, and plant growth at high temperature in Arabidopsis J Exp Bot 68:5981–5981

Chiu RS, Pan S, Zhao R, Gazzarrini S. 2016. ABA-dependent inhibition of the ubiquitin proteasome system during germination at high temperature in Arabidopsis. The Plant Journal 88: 749–761

Curaba J, Moritz T, Blervaque R, Parcy F, Raz V, Herzog M, Vachon G. 2004. AtGA3ox2, a key gene responsible for bioactive gibberellin biosynthesis, is regulated during embryogenesis by LEAFY COTYLEDON2 and FUSCA3 in Arabidopsis. Plant Physiol. 136:3660–9.

Clough SJ, and Bent AF. 1998. Floral dip: a simplified method for Agrobacterium-mediated transformation of Arabidopsis thaliana. Plant J 16:735–743

de Lucas M, Pu L, Turco G, Gaudinier A, Morao AK, Harashima H, Kim D, Ron M, Sugimoto K, Roudier F, et al. 2016. Transcriptional Regulation of Arabidopsis Polycomb Repressive Complex 2 Coordinates Cell-Type Proliferation and Differentiation. Plant Cell 28:2616–2631

de Vries SC, and Weijers D. 2017. Plant embryogenesis. Curr Biol 27:R870–R873

Dean G, Cao Y, Xiang D, Provart NJ, Ramsay L, Ahad A, White R, Selvaraj G, Datla R, and Haughn G. 2011. Analysis of gene expression patterns during seed coat development in Arabidopsis. Mol Plant 4:1074–1091

Deplancke B, Vermeirssen V, Arda HE, Martinez NJ, and Walhout AJ. 2006. Gateway-compatible yeast one-hybrid screens. CSH Protoc 2006:doi: 10.1101/pdb.prot4590

Dresselhaus T, Sprunck S, and Wessel GM. 2016. Fertilization Mechanisms in Flowering Plants. Curr Biol 26:R125–139

Drews GN, and Koltunow AM. 2011. The female gametophyte. Arabidopsis Book 9:e0155

Duong S, Vonapartis E, Li CY, Patel S, and Gazzarrini S. 2017. The E3 ligase ABI3-INTERACTING PROTEIN2 negatively regulates FUSCA3 and plays a role in cotyledon development in Arabidopsis thaliana. J Exp Bot 68:1555–1567

Elliott RC, Betzner AS, Huttner E, Oakes MP, Tucker WQ, Gerentes D, Perez P, and Smyth DR. 1996. AINTEGUMENTA, an APETALA2-like gene of Arabidopsis with pleiotropic roles in ovule development and floral organ growth. Plant Cell 8:155–168

Fatihi A, Boulard C, Bouyer D, Baud S, Dubreucq B, and Lepiniec L. 2016. Deciphering and modifying LAFL transcriptional regulatory network in seed for improving yield and quality of storage compounds. Plant Sci 250:198–204

Figueiredo DD, and Kohler C. 2018. Auxin: a molecular trigger of seed development. Genes Dev 32:479–490

Figueiredo DD, Batista RA, Roszak PJ, and Kohler C. 2015. Auxin production couples endosperm development to fertilization. Nat Plants 1:15184

Figueiredo DD, Batista RA, Roszak PJ, Hennig L, and Kohler C. 2016. Auxin production in the endosperm drives seed coat development in Arabidopsis. eLife 5

Gasser CS, and Skinner DJ. 2019. Development and evolution of the unique ovules of flowering plants. Curr Top Dev Biol 131:373–399

Gazzarrini S, Tsuchiya Y, Lumba S, Okamoto M, and McCourt P. 2004. The transcription factor FUSCA3 controls developmental timing in Arabidopsis through the hormones gibberellin and abscisic acid. Devel Cell 7:373–385

Gietz RD, and Schiestl RH. 2007. Large-scale high-efficiency yeast transformation using the LiAc/SS carrier DNA/PEG method. Nat Protoc 2:38–41

Gregis V, Sessa A, Dorca-Fornell C, and Kater MM. 2009. The Arabidopsis floral meristem identity genes AP1, AGL24 and SVP directly repress class B and C floral homeotic genes. Plant J 60:626–637

Grossniklaus U, Vielle-Calzada JP, Hoeppner MA, and Gagliano WB. 1998. Maternal control of embryogenesis by MEDEA, a polycomb group gene in Arabidopsis. Science 280: 446–450

Hecker A, Brand LH, Peter S, Simoncello N, Kilian J, Harter K, Gaudin V, and Wanke D. 2015. The Arabidopsis GAGA-Binding Factor BASIC PENTACYSTEINE6 Recruits the POLYCOMB-REPRESSIVE COMPLEX1 Component LIKE HETEROCHROMATIN PROTEIN1 to GAGA DNA Motifs. Plant Physiol 168:130–141

Heckman KL, and Pease LR. 2007. Gene splicing and mutagenesis by PCR-driven overlap extension. Nat Protoc 2:924–932

Jia H, Suzuki M, and McCarty DR. 2014. Regulation of the seed to seedling developmental phase transition by the LAFL and VAL transcription factor networks. Wiley Interdiscip Rev Dev Biol 3:135–145

Keith K, Kraml M, Dengler NG, and McCourt P. 1994. fusca3: A Heterochronic Mutation Affecting Late Embryo Development in Arabidopsis. Plant Cell 6:589–600

Kiyosue T, Ohad N, Yadegari R, Hannon M, Dinneny J, Wells D, Katz A, Margossian L, Harada JJ, Goldberg RB, et al. 1999. Control of fertilization-independent endosperm development by the MEDEA polycomb gene in Arabidopsis. Proc Natl Acad Sci USA 96:4186–4191

Klucher KM, Chow H, Reiser L, and Fischer RL. 1996. The AINTEGUMENTA gene of Arabidopsis required for ovule and female gametophyte development is related to the floral homeotic gene APETALA2. Plant Cell 8:137–153

Kohler C, and Grossniklaus U. 2002. Epigenetic inheritance of expression states in plant development: the role of Polycomb group proteins. Curr Opin Cell Biol 14:773–779

Kooiker M, Airoldi CA, Losa A, Manzotti PS, Finzi L, Kater MM, and Colombo L. 2005. BASIC PENTACYSTEINE1, a GA binding protein that induces conformational changes in the regulatory region of the homeotic arabidopsis gene SEEDSTICK. Plant Cell 17:722–729

Kurihara D, Mizuta Y, Sato Y, and Higashiyama T. 2015. ClearSee: a rapid optical clearing reagent for whole-plant fluorescence imaging. Devel 142:4168–4179

Lafon-Placette C, and Kohler C. 2014. Embryo and endosperm, partners in seed development. Curr Opin Plant Biol 17:64–69

Larsson E, Vivian-Smith A, Offringa R, and Sundberg E. 2017. Auxin Homeostasis in Arabidopsis Ovules Is Anther-Dependent at Maturation and Changes Dynamically upon Fertilization. Front Plant Sci doi: 10.3389/fpls.2017.01735

Lau S, Slane D, Herud O, Kong J, and Jurgens G. 2012. Early embryogenesis in flowering plants: setting up the basic body pattern. Annu Rev Plant Biol 63:483–506

Lepiniec L, Devic M, Roscoe TJ, Bouyer D, Zhou DX, Boulard C, Baud S, and Dubreucq B. 2018. Molecular and epigenetic regulations and functions of the LAFL transcriptional regulators that control seed development. Plant Reprod 31:291–307

Liu J, Deng S, Wang H, Ye J, Wu HW, Sun HX, and Chua NH. 2016. CURLY LEAF Regulates Gene Sets Coordinating Seed Size and Lipid Biosynthesis. Plant Physiol 171:424–436

Lotan T, Ohto M, Yee KM, West MAL, Lo R, Kwong RW, Yamagishi K, Fischer RL, Goldberg RB, and Harada JJ. 1998. Arabidopsis LEAFY COTYLEDON1 is sufficient to induce embryo development in vegetative cells. Cell 93:1195–1205

Lu QS, dela Paz J, Pathmanathan A, Chiu RS, Tsai AYL, and Gazzarrini S. 2010. The C-terminal domain of FUSCA3 negatively regulates mRNA and protein levels, and mediates sensitivity to the hormones abscisic acid and gibberellic acid in Arabidopsis. Plant J 64:100–113

Lumba S, Tsuchiya Y, Delmas F, Hezky J, Provart NJ, Shi Lu Q, McCourt P, and Gazzarrini S. 2012. The embryonic leaf identity gene FUSCA3 regulates vegetative phase transitions by negatively modulating ethylene-regulated gene expression in Arabidopsis. BMC Biol doi: 10.1186/1741-7007-10-8

Luo M, Bilodeau P, Dennis ES, Peacock WJ, and Chaudhury A. 2000. Expression and parent-of-origin effects for FIS2, MEA, and FIE in the endosperm and embryo of developing Arabidopsis seeds. Proc Natl Acad Sci USA 97:10637–10642

Makarevich G, Leroy O, Akinci U, Schubert D, Clarenz O, Goodrich J, Grossniklaus U, and Kohler C. 2006. Different Polycomb group complexes regulate common target genes in Arabidopsis. EMBO Rep 7:947–952

Meister RJ, Williams LA, Monfared MM, Gallagher TL, Kraft EA, Nelson CG, and Gasser CS. 2004. Definition and interactions of a positive regulatory element of the Arabidopsis INNER NO OUTER promoter. Plant J 37:426–438

Mitsuda N, Ikeda M, Takada S, Takiguchi Y, Kondou Y, Yoshizumi T, Fujita M, Shinozaki K, Matsui M, and Ohme-Takagi M. 2010. Efficient Yeast One-/Two-Hybrid Screening Using a Library Composed Only of Transcription Factors in Arabidopsis thaliana. Plant Cell Physiol 51:2145–2151

Monfared MM, Simon MK, Meister RJ, Roig-Villanova I, Kooiker M, Colombo L, Fletcher JC, and Gasser CS. 2011. Overlapping and antagonistic activities of BASIC PENTACYSTEINE genes affect a range of developmental processes in Arabidopsis. Plant J 66:1020–1031

Mozgova I, Köhler C, Hennig L. 2015. Keeping the gate closed: functions of the polycomb repressive complex PRC2 in development. Plant J. 83: 121–32.

Mozgova I, Hennig L. 2015. The polycomb group protein regulatory network. Annu Rev Plant Biol. 66: 269–96.

Robert HS. 2019. Molecular Communication for Coordinated Seed and Fruit Development: What Can We Learn from Auxin and Sugars? Int J Mol Sci doi: 10.3390/ijms20040936

Robert HS, Park C, Gutierrez CL, Wojcikowska B, Pencik A, Novak O, Chen JY, Grunewald W, Dresselhaus T, Friml J, et al. 2018. Maternal auxin supply contributes to early embryo patterning in Arabidopsis. Nat Plants 4:548–553

Roscoe TJ, Vaissayre V, Paszkiewicz G, Clavijo F, Kelemen Z, Michaud C, Lepiniec L, Dubreucq B, Zhou DX, and Devic M. 2019. Regulation of FUSCA3 expression during seed development in Arabidopsis. Plant Cell Physiol 60:476–487

Roszak P, and Kohler C. 2011. Polycomb group proteins are required to couple seed coat initiation to fertilization. Proc Natl Acad Sci USA 108:20826–20831

Schmid M, Davison TS, Henz SR, Pape UJ, Demar M, Vingron M, Scholkopf B, Weigel D, and Lohmann JU. 2005. A gene expression map of Arabidopsis thaliana development. Nat Genet 37:501–506

Schneitz K, Hülskamp M, and Pruitt RE. 1995. Wild-type ovule development in Arabidopsis thaliana: a light microscope study of cleared whole-mount tissue. Plant J 7:731–749

Simonini S, and Kater MM. 2014. Class I BASIC PENTACYSTEINE factors regulate HOMEOBOX genes involved in meristem size maintenance. J Exp Bot 65:1455–1465

Simonini S, Roig-Villanova I, Gregis V, Colombo B, Colombo L, and Kater MM. 2012. Basic pentacysteine proteins mediate MADS domain complex binding to the DNA for tissue-specific expression of target genes in Arabidopsis. Plant Cell 24:4163–4172

Sreenivasulu N, and Wobus U. 2013. Seed-Development Programs: A Systems Biology-Based Comparison Between Dicots and Monocots. Annu Rev Plant Biol 64:189–217

Stone SL, Kwong LW, Yee KM, Pelletier J, Lepiniec L, Fischer RL, Goldberg RB, and Harada JJ. 2001. LEAFY COTYLEDON2 encodes a B3 domain transcription factor that induces embryo development. Proc Natl Acad Sci USA 98:11806–11811

Suh MC, Samuels AL, Jetter R, Kunst L, Pollard M, Ohlrogge J, and Beisson F. 2005. Cuticular lipid composition, surface structure, and gene expression in Arabidopsis stem epidermis. Plant Physiol 139:1649–1665

Swanson R, Clark T, and Preuss DJSPR. 2005. Expression profiling of Arabidopsis stigma tissue identifies stigma-specific genes. Sex Plant Reprod 18:163–171

Tsai AY, Gazzarrini S. 2012. AKIN10 and FUSCA3 interact to control lateral organ development and phase transitions in Arabidopsis. The Plant Journal 69: 809–21.

Tsuchiya Y, Nambara E, Naito S, and McCourt P. 2004. The FUS3 transcription factor functions through the epidermal regulator TTG1 during embryogenesis in Arabidopsis. Plant J 37:73–81

Tsuda K, Abraham-Juarez MJ, Maeno A, Dong Z, Aromdee D, Meeley R, Shiroishi T, Nonomura KI, and Hake S. 2017. KNOTTED1 Cofactors, BLH12 and BLH14, Regulate Internode Patterning and Vein Anastomosis in Maize. Plant Cell 29:1105–1118

Vashisht D, and Nodine MD. 2014. MicroRNA functions in plant embryos. Biochem Soc Trans 42:352–357

Zhou Y, Wang Y, Krause K, Yang T, Dongus JA, Zhang Y, Turck F. 2018. Telobox motifs recruit CLF/SWN-PRC2 for H3K27me3 deposition via TRB factors in Arabidopsis. Nat Genet. 50:638–644

Wang D, Tyson MD, Jackson SS, and Yadegari R. 2006. Partially redundant functions of two SET-domain polycomb-group proteins in controlling initiation of seed development in Arabidopsis. Proc Natl Acad Sci USA 103:13244–13249

Wang G, and Kohler C. 2017. Epigenetic processes in flowering plant reproduction. J Exp Bot 68:797–807

Wu J, Wu W, Liang J, Jin Y, Gazzarrini S, He J, and Yi M. 2019. GhTCP19 Transcription Factor Regulates Corm Dormancy Release by Repressing GhNCED Expression in Gladiolus. Plant Cell Physiol 60:52–62

Wu J, Jin Y, Liu C, Vonapartis E, Liang J, Wu W, Gazzarrini S, He J, and Yi M. 2018. GhNAC83 inhibits corm dormancy release by regulating ABA signaling and cytokinin biosynthesis in Gladiolus hybridus. J Exp Bot 70:1221–1237

Xiao J, Jin R, Yu X, Shen M, Wagner JD, Pai A, Song C, Zhuang M, Klasfeld S, He C, et al. 2017. Cis and trans determinants of epigenetic silencing by Polycomb repressive complex 2 in Arabidopsis. Nat Genet 49:1546

