## Supplementary figures for "Spatiotemporal restriction of *FUSCA3* expression by class I BPC promotes ovule development and coordinates embryo and endosperm growth"

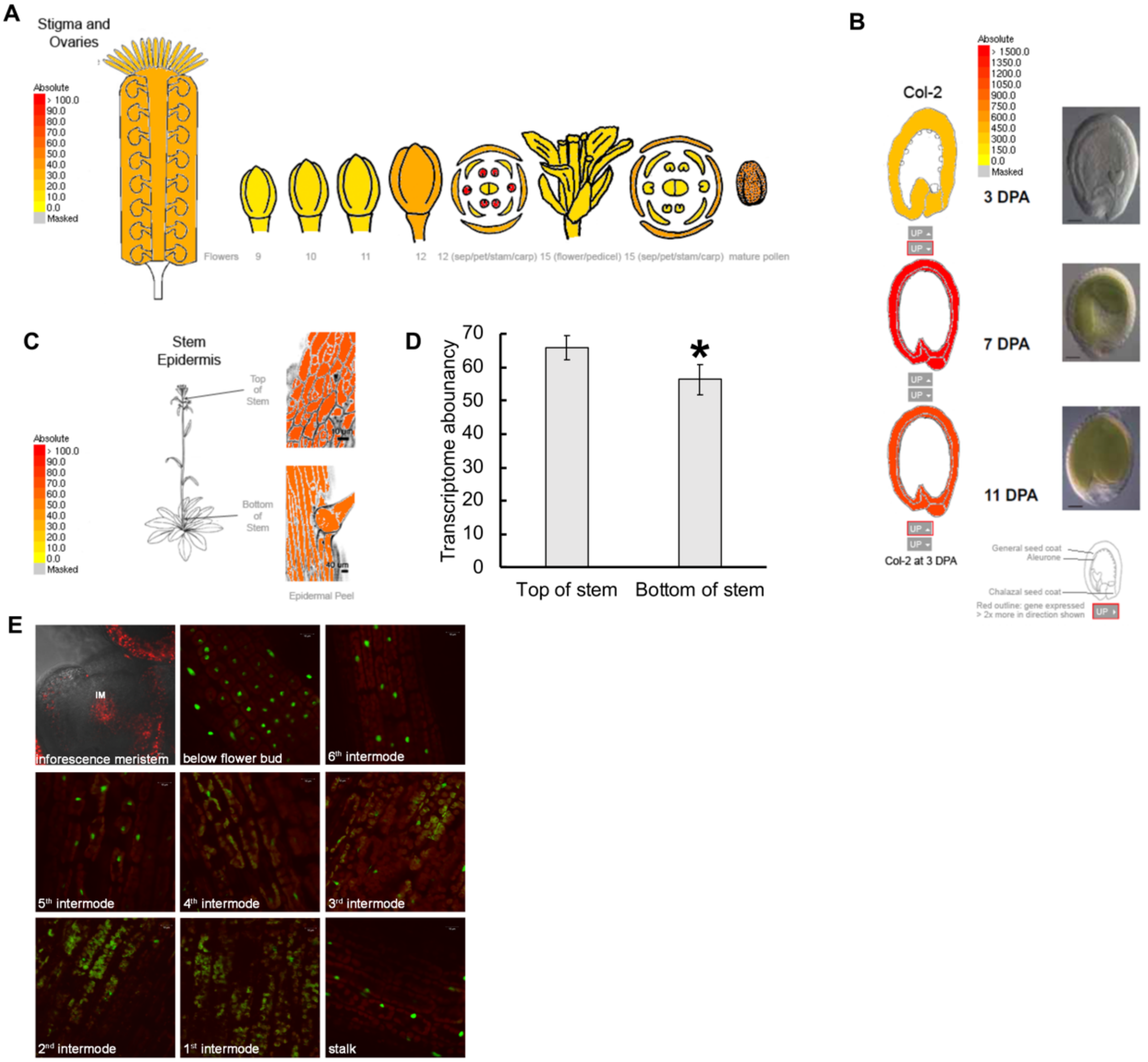


**Supplemental Figure 1 *FUS3* expression profile in reproductive tissues and stem epidermis.** The images were generated by the eFP browser (www.bar.utoronto.ca) using microarray data (Schmid et al., 2005; Suh et al., 2005; Swanson et al., 2005; Dean et al., 2011). **A**, *FUS3* expression profile in the pistil and different stages of flowers. **B**, The expression profile of *FUS3* in the seed coat at the heart, bending and walking embryo stage. **C**, The expression pattern of *FUS3* in the stem epidermis. **D**, *FUS3* transcripts are higher at the top of the inflorescence stem compared to bottom (*: $p<0.05$; student t-test). **E**, *pFUS3:FUS3ΔC-GFP* fluorescence appears stronger in the epidermis of internodes closer to the flower buds and weaker at the bottom of the stem. Identical confocal settings were used.

-615 gaccagtaagtaatgagagttaaattgactttaaaactagagttaaatgtggtagagagagagaagaagacgccattgaattctggaaaaaaggaaagaa
caaacaaaca

-515 gcagaacttttgctacacttgttcaccatgcacaatccactcgtccctttgctcttggttctaaactttaacctaacctaatctctaaactatggttgag

-415 tgttagtttagtggcattagatcttgttgcgttactatcatcataataatcttatcaagatttagaaaatcttaatcttccaaaaccctagattcatgaa

-315 tgtaggttatacaaatgtcacttccttaccataaatagtaaaacaaaaatcattaaaaaccaatggtaaacccaaatctctgattgcccagcgtccattt

-215 taagcctctgtttcgatctgcacaacacacatacacacaacacaacacaaaccacagtctctcttctctctctatctatcttctcttctctctctatc
tgtttg tttgtttg tttg tttgtttg tt

-115 tctatcactgaaacccaaagagatccaccatttgttctttttccttcacacagagaactgttttcttccacacttccttttactaggcagtgttaacc
tg

-15 aattgagagagaaaa -1
caaacaaa

**Supplemental Figure 2 Location of (GA/CT)$_n$ mutated in the *pFUS3* genomic region used in Y1H.** The (GA/CT)$_n$ motifs that were mutated in the *FUS3* sequence upstream of the start codon (base pairs -1 to -615) are highlighted in yellow. Mutated G/C residues are marked in red. Telo boxes binding sites of AZF1 (Xiao et al., 2017) are marked in blue. The repressive region (126bp) that associated with MEA and H3K27me3 repressive marks (Makarevich et al., 2006), and was deleted in promoter studies (Roscoe et al. (2019) is underlined.

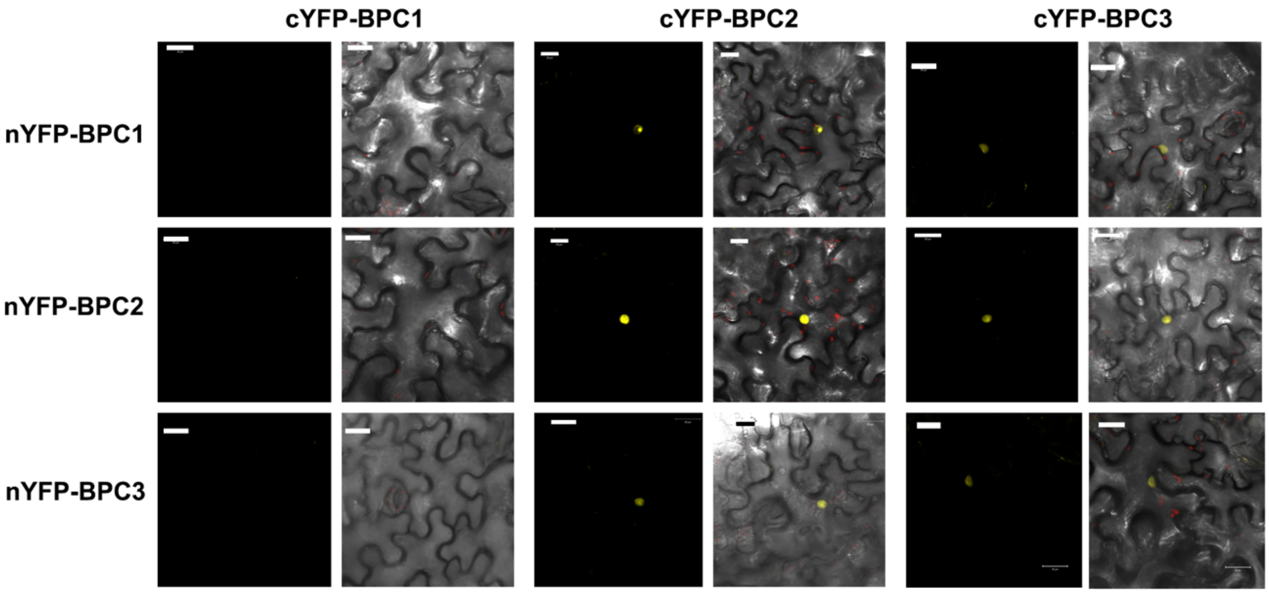


**Supplemental Figure 3 Class I BPC family members form homo- or hetero- dimers.** BPC1, BPC2 and BPC3 form heterodimers with each other. Only BPC2 and BPC3 form homodimers. Lack of interaction between FUS3 and BPCs or FIS-PRC2 in BiFC assays is shown as the negative control in Supplemental Figure 5.

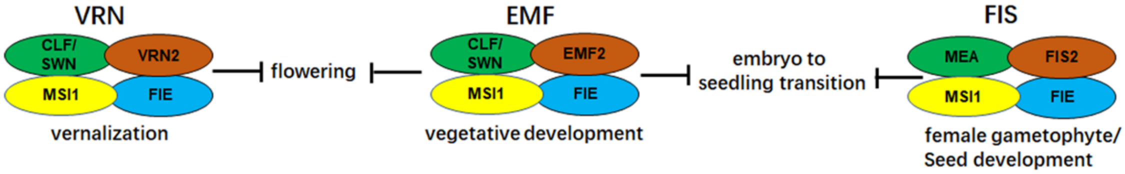


**Supplemental Figure 4 PRC2 complexes in Arabidopsis.** There are three different types of PRC2 in Arabidopsis depending on subunit composition - VRN-PRC2, EMF-PRC2 and FIS-PRC2 -, which regulate vernalization, vegetative development and female gametophyte/seed development, respectively (Mozgova et al., 2015).

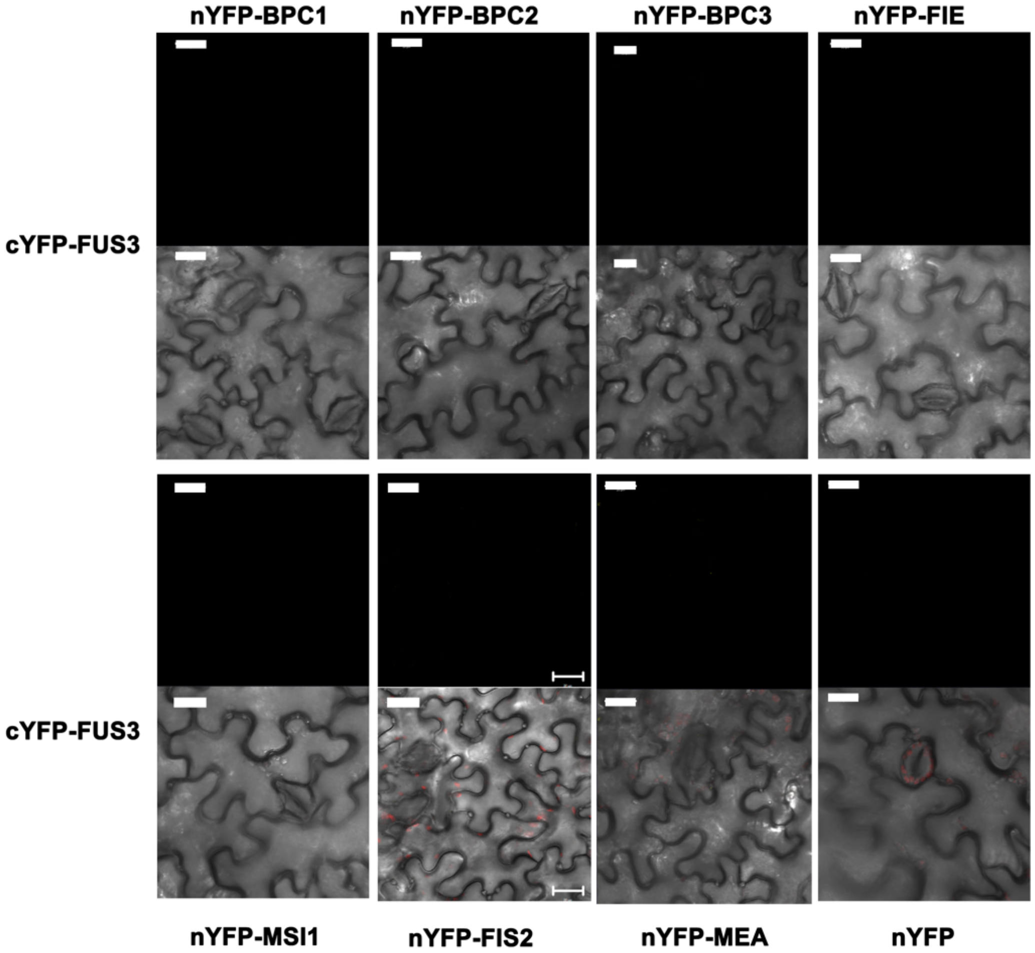


**Supplemental Figure 5 FUS3 protein does not interact with FIS-PRC2 complex and Class I BPC family members using BiFC.** Negative control showing that FUS3 does not interact with FIE, MSI1, MEA, FIS2, BPC1, BPC2 or BPC3 in *N. benthamiana* by BiFC.

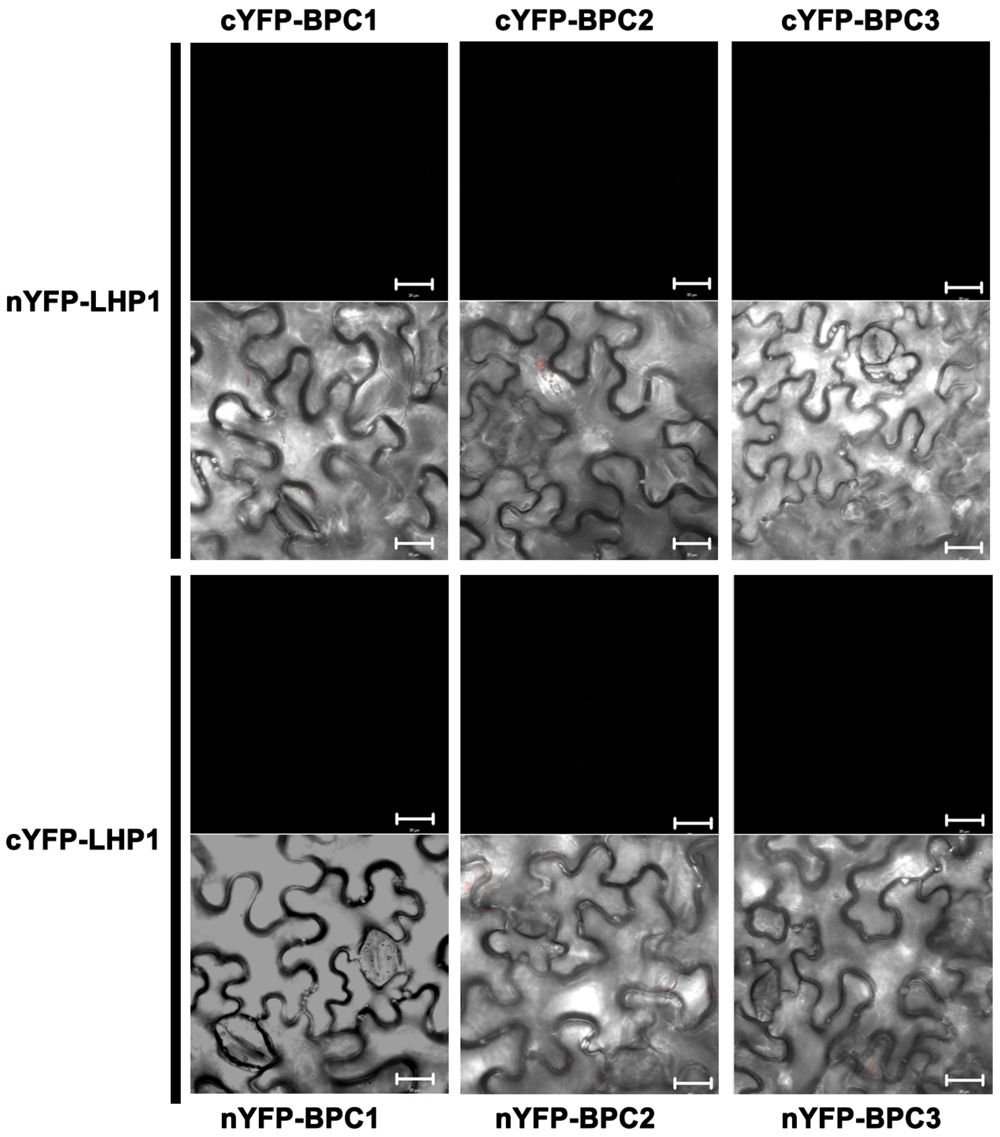


**Supplemental Figure 6 Class I BPC proteins do not interact with LHP1 using BiFC.** Class I BPC proteins do not interact with LHP1 in *N. benthamiana* by BiFC.

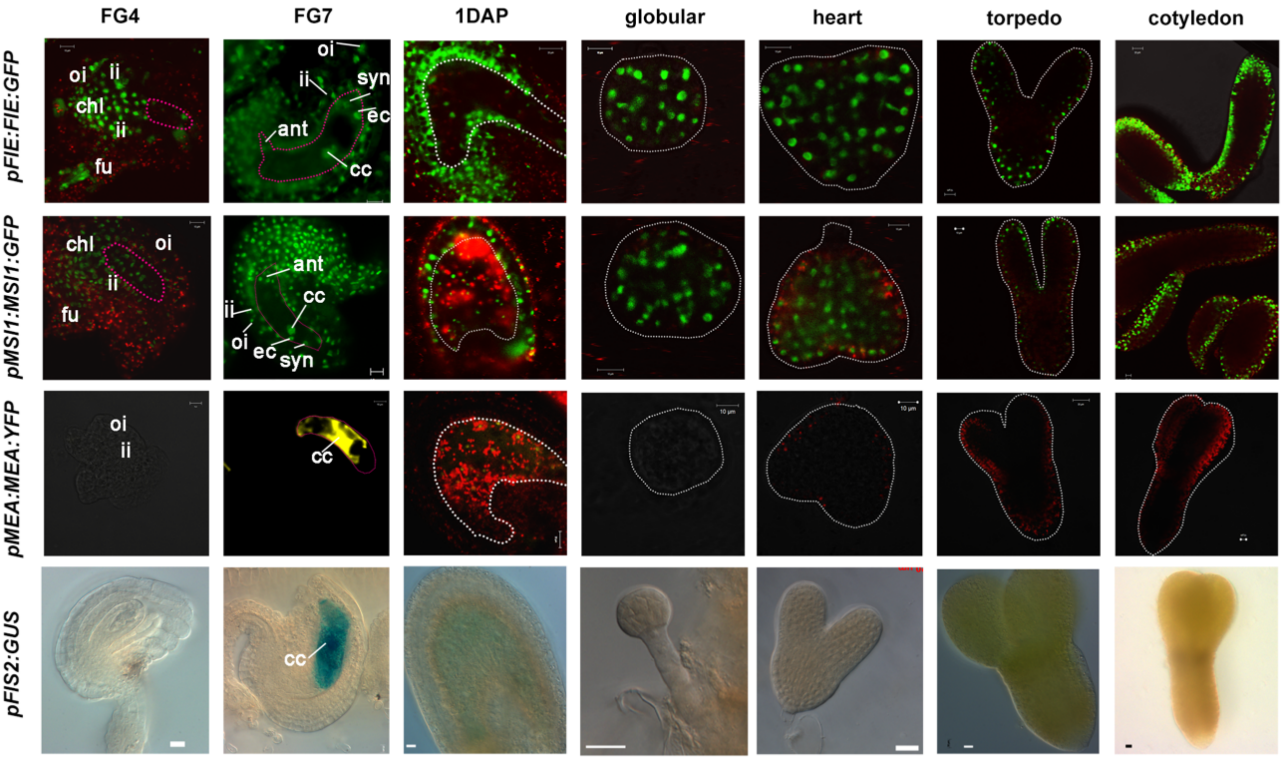


**Supplemental Figure 7 Expression patterns of FIS-PRC2 in ovules and embryos.**

FIS2 complex (FIE, MSI1, MEA and FIS2) expression/localization patterns in ovules before fertilization (FG4 and FG7), 1DAP seeds and embyos at globular, heart, torpedo and cotyledon stages. Ant: antipodals; cc: central cell; chl: chalaza; cze: chalazal endosperm; ec: egg cell; fu: funiculus; ii: inner teguments; mce: micropilar endosperm; oi: outer teguments; pen: peripheral endosperm; pge: pre-globular embryo; sdc: seed coat; sus: suspensor; syn: synergids cell. Pink dashed lines represent the outline of the embryo sac. The ferro/ferricyanide used in GUS staining buffer was 2mM for *pFIS2:GUS*.

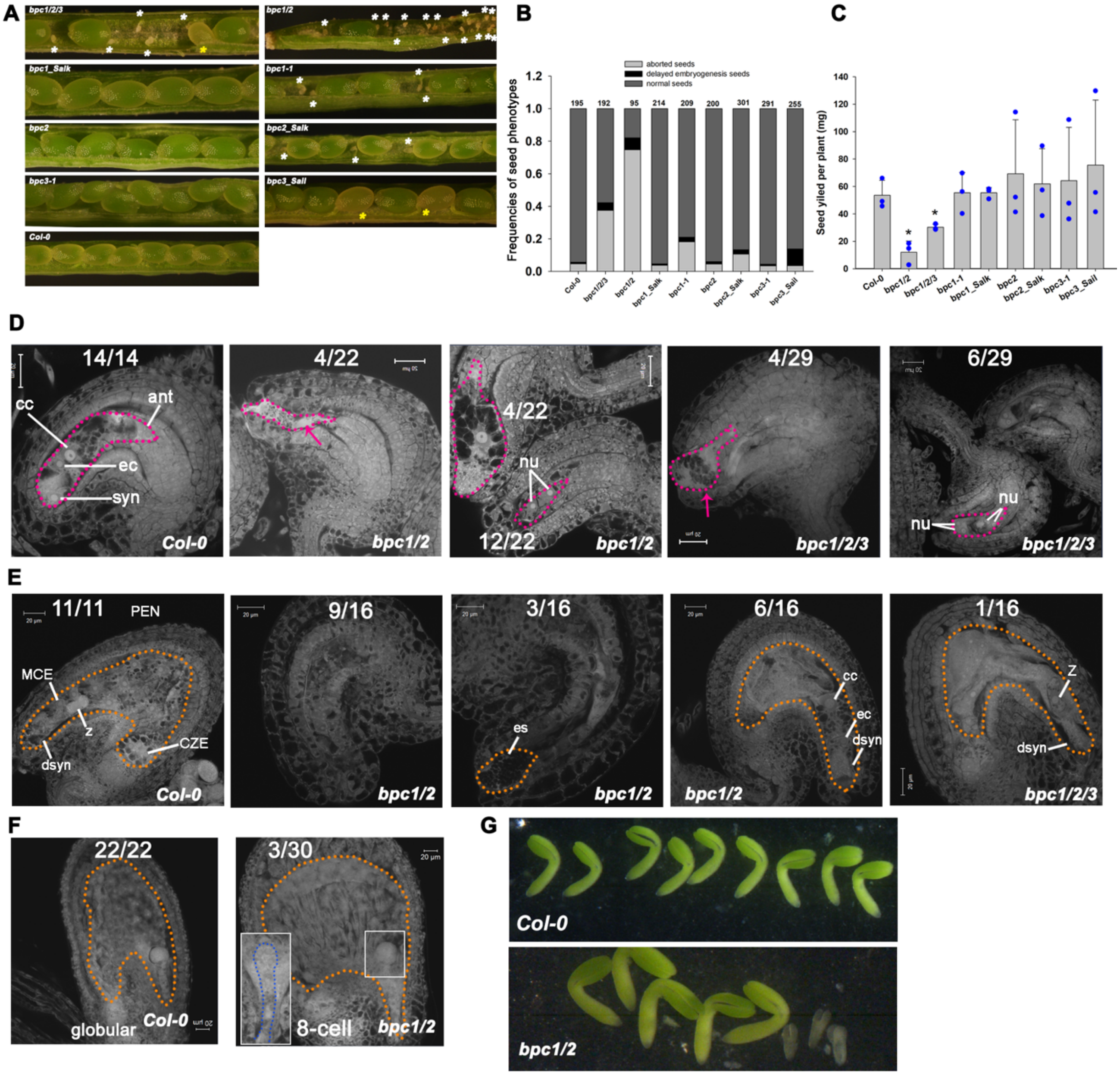


**Supplemental Figure 8 Class I *bpc* mutants show delayed megagametogenesis, seed abortion and delayed embryogenesis. A**, Class I BPC mutants show seed abortion and delayed embryogenesis phenotypes. The white asterisks indicate aborted seeds; the yellow asterisks represent delayed embryogenesis seeds. **B**, The frequencies of seed phenotypes in 10 peeled half-side of *bpc* mutants siliques. Three biological repeats were performed and one representative result is shown. **C**, Seed yield of WT and *bpc* mutants. The error bars represent the SD of three biological replicates (*: $p<0.05$; student t-test was used) **D,** FS12 ovules of Class I BPC mutants showing aborted embryo sac and delayed megagametyogenesis. **E**, Class I BPC mutants show condensed endosperm and unfertilized egg cell with the degenerated synergid cell at 2DAF. **F**, Arrested embryos in *bpc1/2* with an enlarged seed at 3DAP. **G**, At mature stage (11DAP), some *bpc1/2* embryos were arrested at torpedo stage. The pink arrow points to the aborted embryo sac. Ant: antipodals; cc: central cell; cze: chalazal endosperm; ec: egg cell; es: embryo sac; mce: micropilar endosperm; nu: nuclei; pen: peripheral endosperm; dsyn: degenerated synergid cell; z: zygote. Pink dashed lines outline the embryo sac at FS12; the yellow dash lines outline the embryo sac at 2DAF.

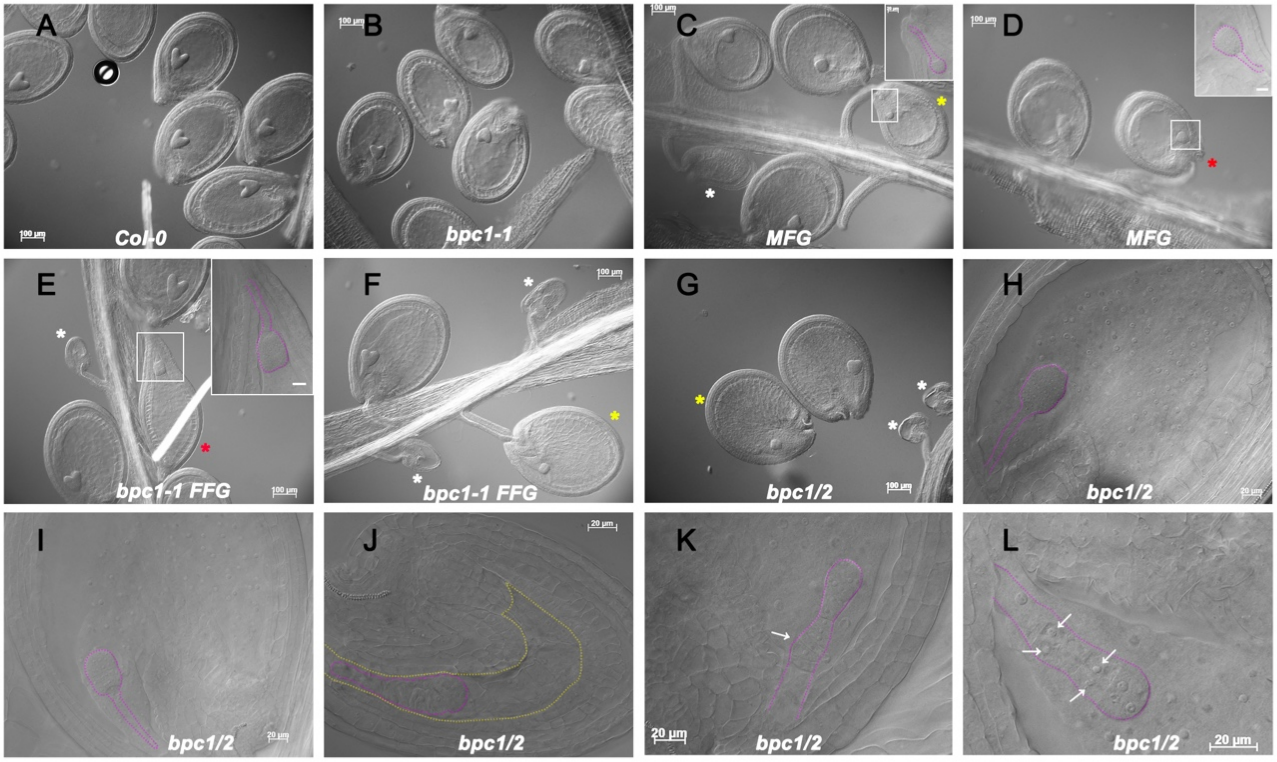


**Supplemental Figure 9 Overexpression of *FUS3* results in embryo defect and over-proliferation of the endosperm nuclei.** Seeds of (**A**) Wildtype, (**B**) *bpc1-1*, (**C, D**) *MFG*, (**E, F**) *bpc1-1 FFG* and (**G-L**) *bpc1/2* at 3 DAP. Pink dashed lines represent the outline of the embryo. The yellow dashed line represents the embryo sac. White arrows indicate the abnormal suspensors. White, yellow or red asterisk indicates the aborted seed, arrested embryo or defective embryo, respectively.

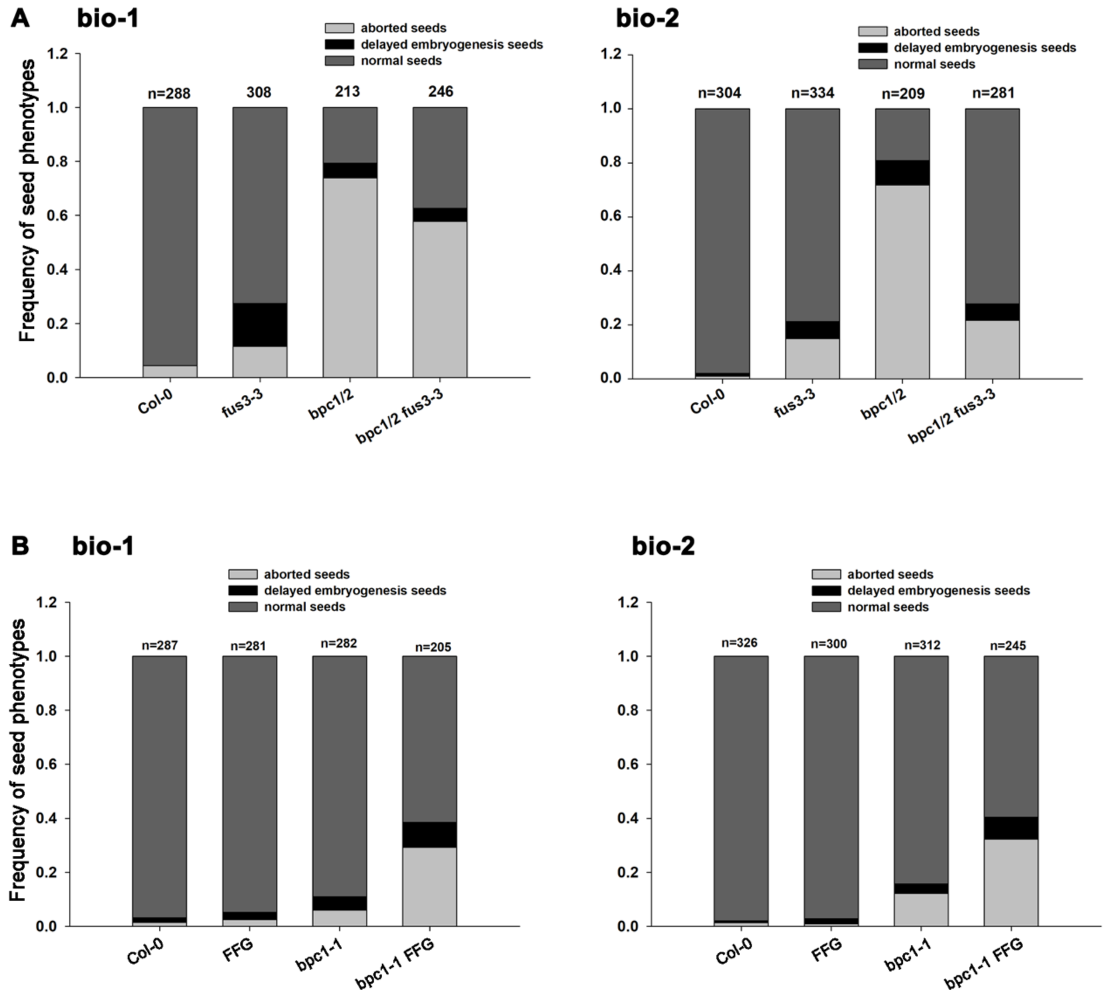


**Supplemental Figure 10 Frequencies of seed phenotypes**. **A, B** The total number of seeds displaying various phenotypes was calculated in 10 peeled siliques (half side) of **A,** WT, *bpc1/2* and *bpc1/2 fus3* mutants, and **B**, WT, *bpc1-1* and *bpc1-1 FFG* mutants. Three biological repeats were performed, and two are shown here. See also Figures 6 and 8.

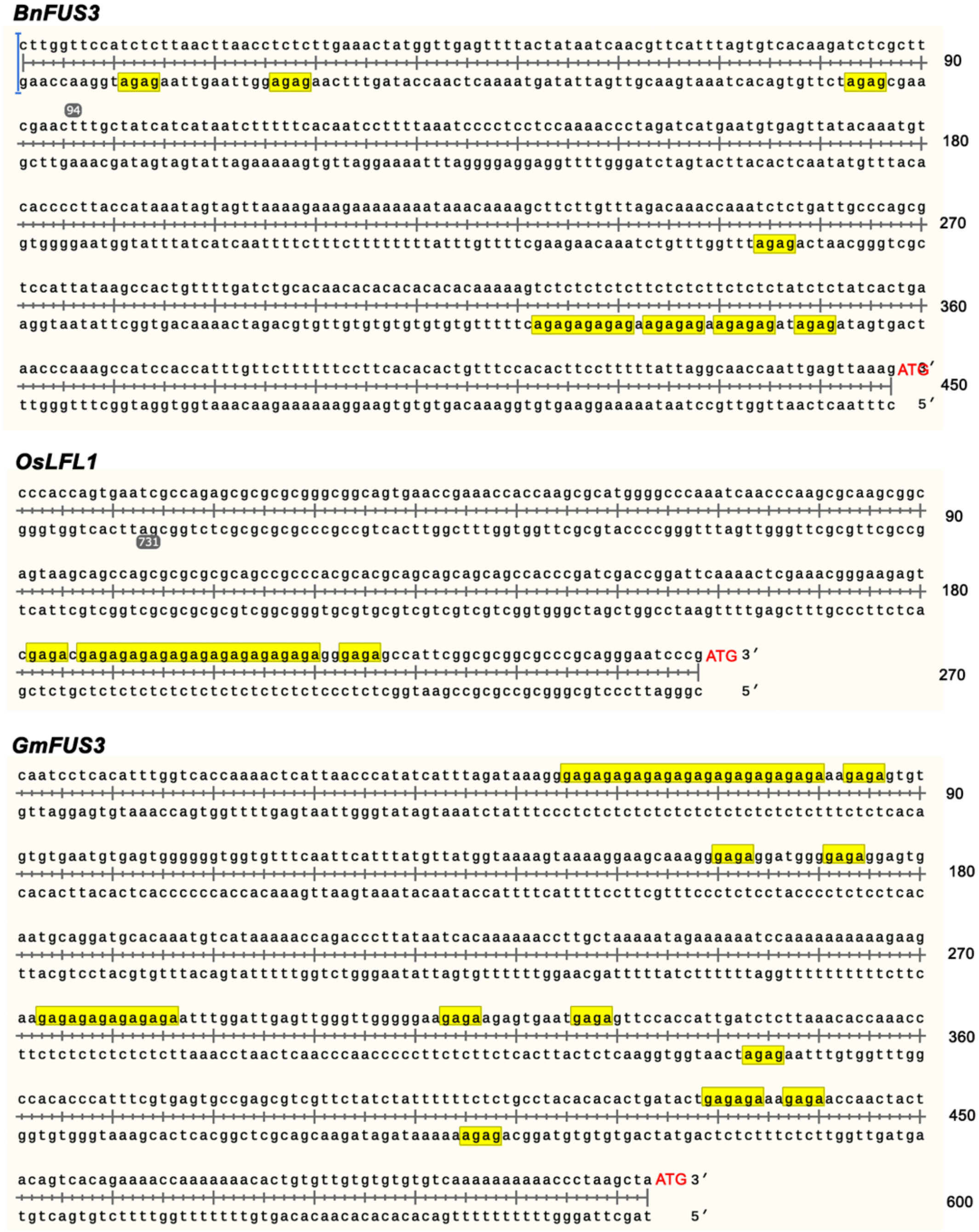


**Supplemental Figure 11 Conserved $(GA/CT)_n$ motifs in orthologous *FUS3* genes.**

Accession number of *FUS3* orthologous genes: *OsLFL1* (LEC2 and FUSCA3-like protein 1**;** GenBank: EF521182.1) from *Oryza sativa*, *BnFUS3* (NCBI: XM_013792060) from *Brassica napus*, *GmFUS3* (Gene ID: LOC100813055) from *Glycine max*.
